# Mechanistic Insights into Proteomic Mutation-Phenotype Linkages from Tiling Mutagenesis Screens

**DOI:** 10.1101/2025.04.17.649336

**Authors:** Wei He, Jen-Wei Huang, Yalong Wang, Samuel B Hayward, Giuseppe Leuzzi, Rongjie Fu, Shuyue Wang, Alina Vaitsiankova, Mark T Bedford, Raphael Guerois, Alberto Ciccia, Han Xu

## Abstract

High-throughput mutagenesis screens are powerful tools for mapping mutations to phenotypes. However, deciphering the molecular mechanisms that link mutations to phenotypic outcomes remains a significant challenge. Here, we present ProTiler-Mut, a versatile computational framework that harnesses tiling mutagenesis screens, which introduce variants across entire protein sequences, to facilitate investigation of mutation-to-phenotype associations at multiple levels, including individual residues, protein substructures, and protein-protein interactions (PPIs). As demonstrated through our analyses of base editing (BE) screens targeting DNA Damage Response (DDR) proteins and T cell regulators, ProTiler-Mut provides novel insights into the mutation-phenotype linkages, including: i) refined classification of mutation that reveals separation-of-function (SOF) category beyond the conventional binary classification of loss-of-function (LOF) and gain-of-function (GOF); ii) definition of phenotype-associated hotspot substructures that enable the inference of the function of unscreened pathogenic mutations; and iii) identification of phenotype-associated PPIs disrupted by functional mutations. Through ProTiler-Mut analyses, we identified a substructure harboring pathogenic GOF mutations that disrupt interactions between the kinases MAPK1 and RSK1, leading to MAPK1 activation and elevated expression of the immune checkpoint receptor PD-1. Furthermore, we demonstrate the applicability of ProTiler-Mut to various mutagenesis screening platforms, highlighting its broad utility and generalizability.

## INTRODUCTION

Understanding protein function is fundamental to molecular biology, with far-reaching implications for deciphering cellular mechanisms and advancing disease treatments. In recent years, CRISPR-based high-throughput screens have transformed functional genomics by enabling the study of gene perturbations at an unprecedented scale^1–4^. To achieve a more granular dissection of protein function, CRISPR knockout (CRISPR-KO) screens that employ tiling sgRNA designs have been developed, where a number of sgRNAs are designed to span entire coding regions of proteins. This strategy facilitates in-depth analyses of discrete protein domains and their individual contributions to protein activity^5–8^. However, CRISPR-KO screens typically introduce small insertions and deletions (indels) and rely on creating double-strand breaks, lowering the resolution of functional analyses and increasing susceptibility to confounding factors^9, 10^.

In early efforts to systematically dissect protein function at single-residue resolution, deep mutational scanning (DMS) enabled the parallel assessment of thousands of amino acid variants^11–13^. Because DMS typically relies on exogenous expression systems, it often fails to reflect the native biological context of the endogenous genomic loci. Subsequently, mutagenesis screens employing CRISPR-based approaches such as base editing (BE) and Homology-directed repair (HDR)-mediated saturation genome editing (SGE), emerged to facilitate *in situ* measurements of mutational effects within the native genomic environment^14–23^. More recently, prime editing (PE) tiling screens have further expanded the scope of possible genetic modifications, allowing targeted insertions, deletions, and substitutions^24–26^.

Despite these technological advances, challenges persist in the interpretation of mutagenesis screening data. First, the sensitivity of mutation-to-phenotype associations is compromised by inherent system biases such as the selection of base editors for mutagenesis, experimental variability in editing efficiency, and limited statistical power for single-mutation analyses. Second, it remains difficult to characterize phenotype-associated mutations and elucidate their functions in the context of protein stability, structure, modifications, and protein-protein interaction (PPI). Third, the sheer volume of functional mutations identified in high-density screens makes manual interpretation impractical, underscoring the need for specialized computational tools that integrate, automate, and streamline the characterization, visualization, and interpretation of identified mutations.

To address these challenges, we developed ProTiler-Mut, a computational framework designed to elucidate the linkages between protein mutations and phenotypic outcomes. By leveraging the enriched and nearly unbiased information generated from tiling mutagenesis screens and integrating experimentally determined or AI-predicted protein structures, ProTiler-Mut enables a robust and comprehensive multi-level analysis of mutation-phenotype relationships, including individual variants, protein substructures, and protein-protein interactions (PPIs). In this paper, we showcase the utility of ProTiler-Mut through two BE-based applications focused on DNA Damage Response (DDR) proteins and anti-tumor T cell regulators. Furthermore, we extend its application to tiling PE, DMS and HDR-mediated SGE screens, demonstrating its broader applicability and generalizability.

## RESULTS

### An overview of tiling mutagenesis screens and ProTiler-Mut

In a tiling mutagenesis screen, mutations are systematically introduced across the entire protein-coding region of target proteins, providing an unbiased, single-residue resolution approach to mutagenesis. The functional impact of individual mutations is then assessed by measuring cellular or molecular phenotypes. For example, in a typical viability screen, fitness-associated mutations result in either enrichment or depletion of mutant cells relative to those carrying neutral mutations. To facilitate high-throughput screening, base editing (BE) or prime editing (PE) technologies are paired with large-scale sgRNA or pegRNA libraries designed for tiled targeting. Alternatively, deep mutational scanning (DMS) employs exhaustive variant libraries transformed into cells, generating a heterogeneous population with diverse mutations. Despite differences in methodology, all tiling mutagenesis screens generate phenotypic profiles for each targeted protein to capture the functional consequences of mutations across the entire protein sequences (Figure 1). Some datasets incorporate phenotypic profiles measured under multiple conditions, such as drug treatments^15, 23, 26^ or cell sorting based on distinct surface markers^13, 27, 28^, expanding the functional insights gained from these screens.

**Figure 1.**
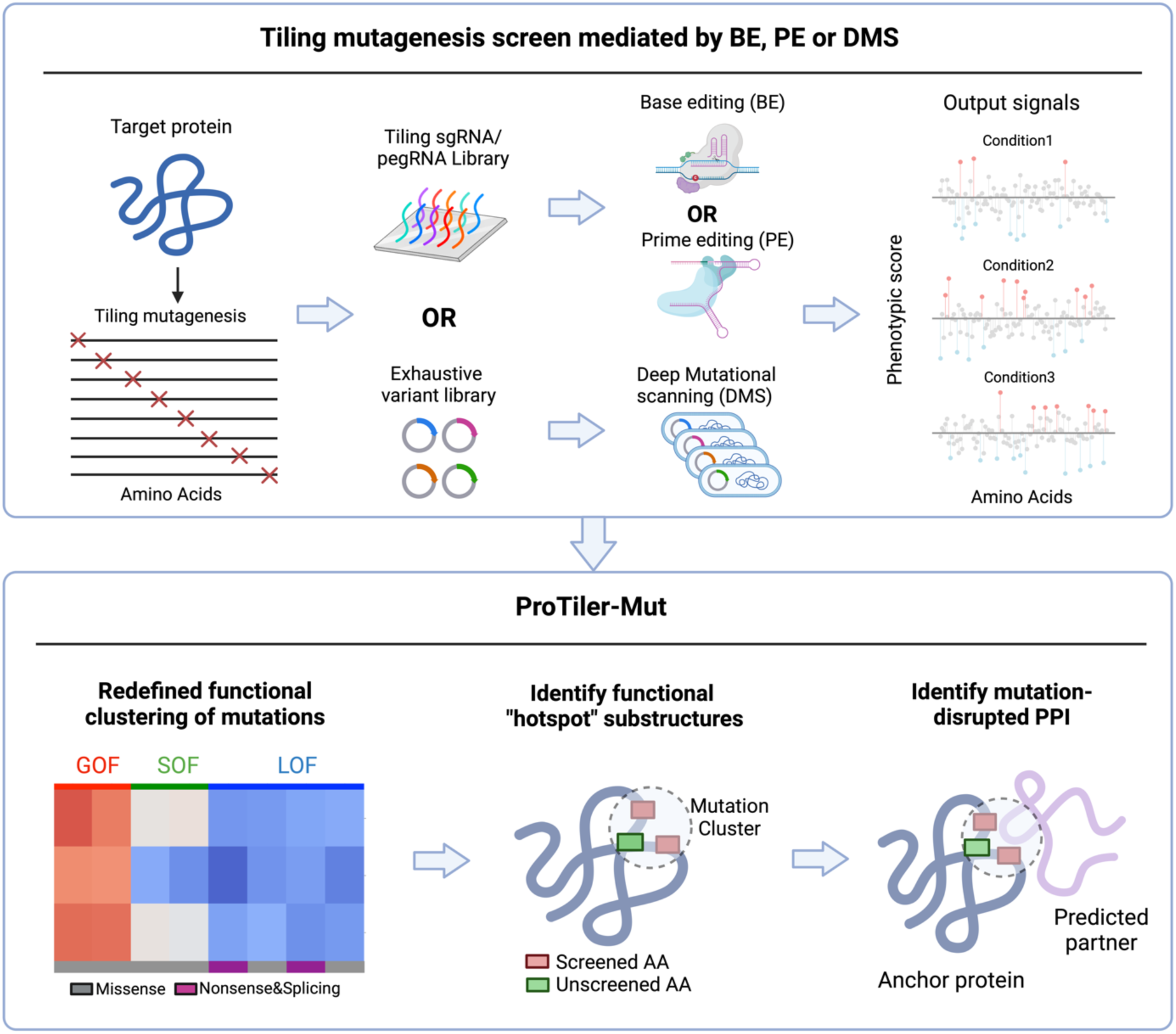
A schematic view of tiling mutagenesis screens and ProTiler-Mut functional modules. The upper panel conceptually represents tiling mutagenesis screening procedures using base editing (BE), prime editing (PE), or deep mutational scanning (DMS), which systematically profile phenotypic scores across the entire protein sequence. ProTiler-Mut leverages these phenotypic profiles to: (i) define functional mutation clusters, (ii) identify “hotspot” substructures that infer the functions of unscreened pathogenic mutations, and (iii) detect mutation-disrupted PPIs (bottom panel).

ProTiler-Mut analyzes phenotypic profiles of mutagenesis for understanding mutation-to-phenotype associations through three core modules (Figure 1). First, it clusters putative functional mutations based on phenotypic similarity and uses nonsense or splicing mutations as references to classify missense mutations as loss-of-function (LOF) or gain-of-function (GOF). Notably, when multiple conditions are considered, unsupervised clustering can reveal categories beyond this binary classification, such as separation of function (SOF) mutations that selectively disrupt specific aspects of a multifunctional protein without eliminating its overall function^15, 29, 30^. Second, ProTiler-Mut identifies 3D functional “hotspot” substructures enriched with mutations that share similar phenotypic outcomes. The de novo mapping of substructures links known protein domains to specific phenotypes and uncovers novel phenotype-associated structures that may involve long-range residue interplays, extending beyond traditional 1D sequence-based domain definitions. These functional substructures also help infer the effects of pathogenic mutations that are not included or are poorly represented in the screen. Finally, ProTiler-Mut leverages high-confidence functional mutations or substructures as “anchors” to investigate potentially disrupted protein-protein interactions (PPIs). This approach narrows the search space for PPIs when combined with AI-predicted interaction structures, pinpointing phenotype-associated interactions with greater confidence for further experimental validation.

### Refined classification of missense mutations beyond binary definition of LOF and GOF

Previous analyses of high-throughput mutagenesis screen data have primarily focused on identifying functional mutations based on the magnitude of phenotypic changes induced by each mutation. While these single-mutation analyses effectively pinpoint the impact of individual mutations, they often lack statistical power and fail to capture the broader functional landscape of mutations. To gain a more comprehensive view of functional patterns across diverse mutations and their inter-relationships, we identify putatively functional mutations using a relatively relaxed threshold (Z score > 2 in at least one condition, see Methods) and group these mutations through hierarchical clustering based on their phenotypic profiles across all conditions.

Applying this exploratory method to a BE screen of 86 DNA Damage Response (DDR) proteins treated with four DNA-damaging agents: cisplatin (CISP), olaparib (OLAP), doxorubicin (DOX) and camptothecin (CPT)^15^, we observed that mutations within the same DDR protein often exhibit more than two distinct phenotypic patterns, surpassing the conventional binary classification of LOF and GOF. For example, as shown in Figure 2a, mutations in the DNA repair factor ERCC2 were grouped into three clusters. Cluster 3 mutations, including all nonsense and splicing mutations, suppress cell growth across all treatment conditions and are indicative of complete LOF. Conversely, mutations in Cluster 1 promote cell growth, representing GOF mutations. A distinct group of Cluster 2 displayed an SOF-like phenotype, where growth suppression occurred selectively under CISP treatment but not under other conditions. Principal component analysis (PCA) further supports that the formation of Cluster 2 (green) is unlikely to be due to random noise (Fig. 2b). Interestingly, mutations in Cluster 2 are enriched at the intra-protein interface between the DEAD domain and the helicase domain (Fig. 2c), suggesting that the SOF phenotype may result from disruption of domain-domain interactions within ERCC2.

**Figure 2.**
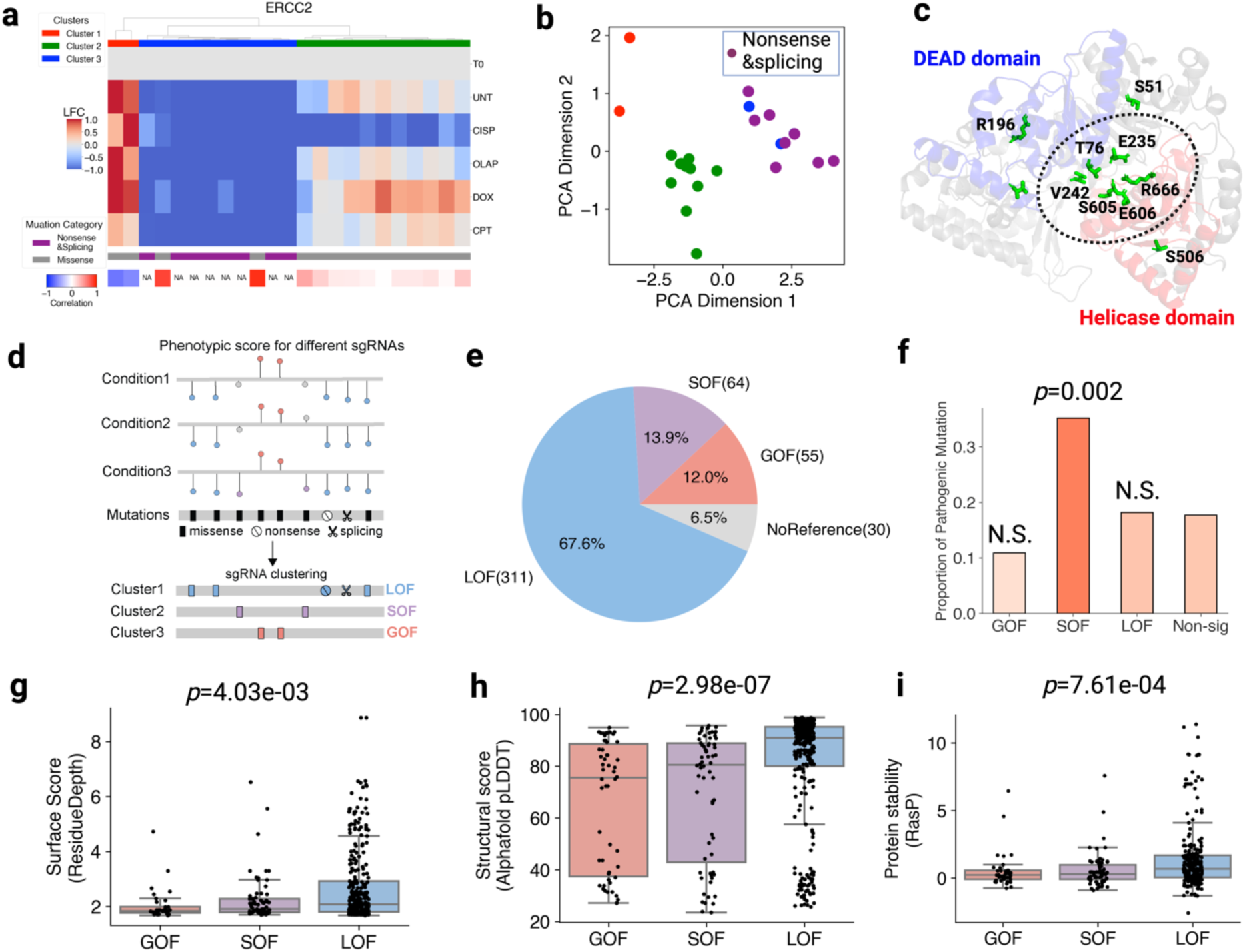
Functional classification of mutations into LOF, SOF, and GOF. **(a)** A heatmap displaying three clusters of functional mutations in ERCC2, categorized based on their phenotypic effects under different drug treatment conditions (UNT: Untreated, CISP: Cisplatin, OLAP: Olaparib, DOX: Doxorubicin, CPT: Camptothecin). Data retrieved from published tiling BE screen data on DNA Damage Response (DDR) proteins^15^. The phenotypic scores represent log fold-change (LFC) in cell viability relative to T0 (pre-selection). Color bars below the heatmap denote mutation types (purple for reference mutations, including nonsense and splicing; gray for missense) and indicate the correlation between missense mutation phenotypic patterns and the averaged profiles of reference mutations. **(b)** Principal component analysis (PCA) of functional mutation profiles of ERCC2. Each dot represents a mutation. Missense mutations are colored according to cluster assignment in (a). Nonsense and splicing mutations are colored in purple. **(c)** Alphafold-predicted structural model of ERCC2, with SOF mutation group (cluster2 in panel a) shown in green sticks. The mutations enriched at the interface between DEAD domain (blue) and Helicase domain (red) are highlighted with a dashed circle. **(d)** A schematic illustration showing classification of LOF, SOF, and GOF mutations, using nonsense and splicing mutations as references for LOF effects. **(e)** Pie chart showing the distribution of mutation categories in 86 DDR proteins. **(f)** Bar chart showing the proportion of pathogenic mutations in GOF, SOF, LOF categories, compared to mutations not associated with significant phenotype in the screen (Non-sig). P-values were calculated using Fisher exact test (N.S.: not significant). **(g-i)** Box plots comparing **(g)** Surface score (the Residue Depth); **(h)** structural scores (AlphaFold pLDDT) and **(i)** free energy change (RasP score) among GOF, SOF, and LOF categories. P-values were calculated using the Kruskal–Wallis test, a non-parametric alternative to one-way ANOVA.

Building on insights from unsupervised mutation clustering, ProTiler-Mut classifies functional missense mutations into GOF, LOF, and SOF categories based on their phenotypic correlation to reference mutations including nonsense and splicing mutations (Fig. 2d). In the DDR dataset, we identified 311 (67.6%) LOF mutations, 64 (13.9%) SOF mutations, and 55 (12.0%) GOF mutations, while 30 (6.5%) remained uncategorized due to the absence of reference mutations (Fig. 2e, Supplementary Table S1). The predominance of LOF mutations aligns with a recent report indicating that 60% of functional missense mutations reduce protein stability^31^. Interestingly, unlike GOF and LOF mutations, SOF mutations showed a significant enrichment of pathogenic variants in the ClinVar dataset (p=0.002, Fig. 2f). Unlike LOF mutations, which generally abolish overall protein activity, SOF mutations tend to selectively disrupt specific functions while preserving others within the same protein. This nuanced impact on DDR proteins can lead to context-dependent pathological effects, highlighting SOF mutations as potential targets for therapeutic intervention.

We further examined the relationship between mutation types and amino acid features. GOF mutations generally exhibit lower residue depth and higher accessible surface area (ASA) values, suggesting their tendency to alter protein interactions at the surface. In contrast, LOF mutations are frequently localized within the hydrophobic core and disrupt overall protein structure (Fig. 2g, Fig. S1a). Additionally, LOF mutations are more prevalent in well-structured and conserved regions (Fig. 2h, Fig. S1b). These distinctions were further supported by ΔΔG predictions using RasP^32^ and DDGun^33^, which highlighted differences in the impact of mutations on protein stability and dynamics (Fig. 2i, Fig. S1c). Notably, SOF mutations exhibit intermediate amino acid feature distributions between the characteristics of GOF and LOF mutations.

We next applied ProTiler-Mut to another BE screen dataset encompassing 385 anti-tumor T cell regulators, where stimulated primary T cells were sorted according to TNF-α, IFN-γ, CD25 or PD-1 levels^28^ (Fig. S2). This analysis identified 6,414 functional mutations, including 5,341 (83.3%) LOF, 267 (4.2%) GOF, 220 (3.4%) SOF, and 586 (9.1%) uncategorized mutations (Fig. S2a, Supplementary Table S2). Despite the predominance of LOF mutations, SOF mutations exhibited the highest enrichment of pathogenic variants, consistent with observations from the DDR dataset (p=0.03, Fig. S2b). Furthermore, the LOF, SOF, and GOF mutations identified in T cell regulators recapitulated the distribution patterns of surface score, conservation, and protein stability observed in DDR proteins (Fig. S2c-e). These findings indicate that ProTiler-Mut provides a biologically meaningful and reproducible classification of missense mutations, whereas the proportion of each mutation category may vary depending on the functional roles of the targeted proteins.

### Hotspot substructures define functional units in 3D protein structure and link unscreened pathogenic mutations to phenotypic patterns

As illustrated in Figure 2c, mutations with similar phenotypic patterns often cluster in close spatial proximity within a protein’s 3D structure. Building on this observation, we developed 3D-RRA (Robust Rank Aggregation), an extension of our previous RRA algorithm^34^, to identify hotspot substructures enriched with functional mutations sharing similar phenotypic patterns (Fig. 3a; see Methods for details). This method increases statistical power by integrating multiple mutations rather than relying on single-mutation analysis. Importantly, this approach can be applied broadly to any protein, leveraging the extensive availability of monomeric 3D structures predicted by AlphaFold^35, 36^.

**Figure 3.**
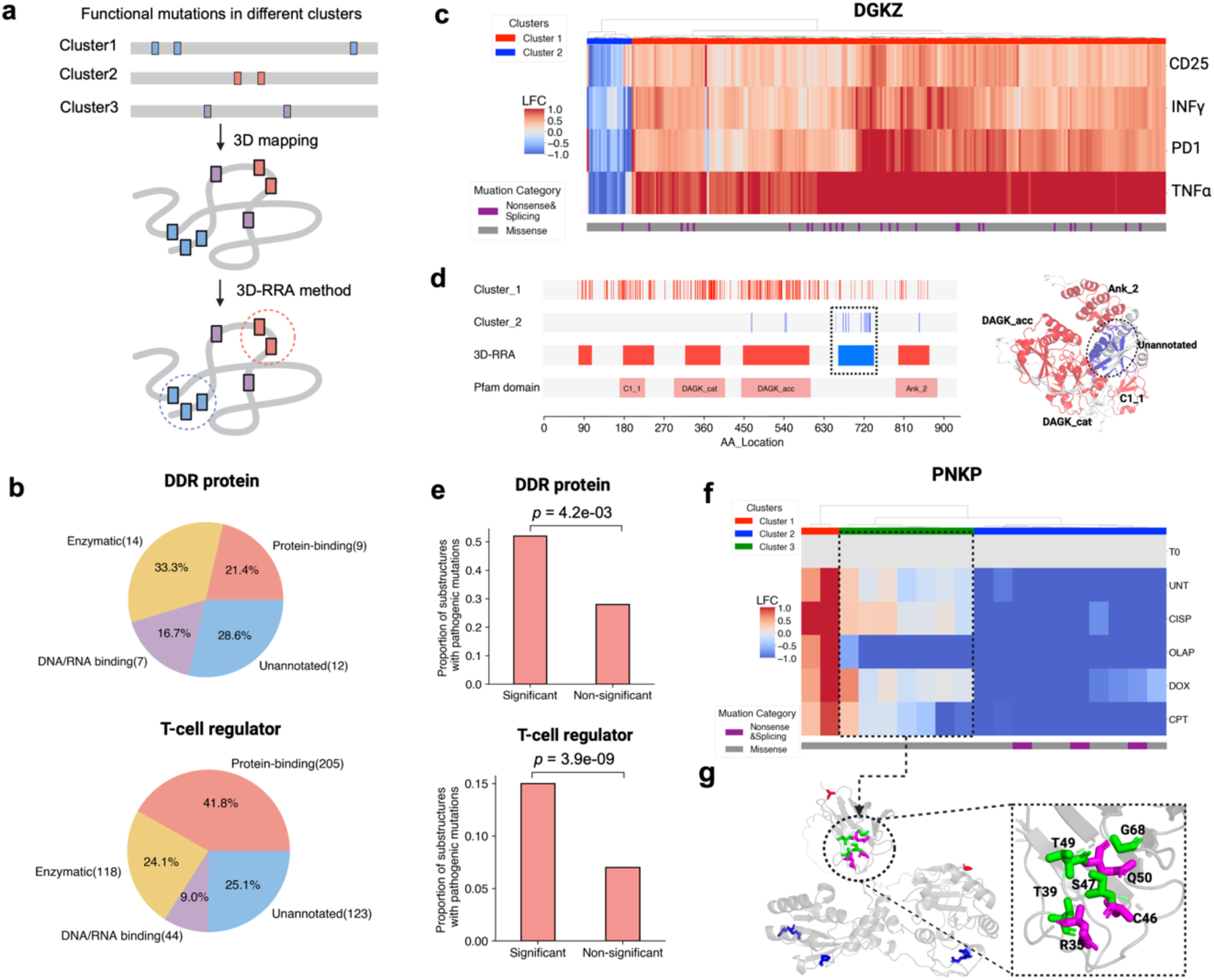
ProTiler-Mut identifies functional “hotspot” substructures. **(a)** Schematic illustration of the workflow for identifying hotspot substructures. **(b)** Pie charts showing the distribution of Pfam protein domain types overlapping the substructures in DDR proteins (upper) and the T-cell regulators (lower). **(c)** Heatmap showing two clusters (LOF, red; GOF, blue) of functional mutations in DGKZ. Data retrieved from published BE screen data on T-cell regulators^28^, where cells harboring mutations were sorted with cytokine secretion (TNF-α, INF-γ) or surface marker expressions (CD25, PD-1). The bottom color bar indicates mutation types (nonsense/splicing in purple, missense in gray). **(d)** The left panel shows the distribution of LOF (red) and GOF (blue) mutations in (c) across the 1D protein sequence of DGKZ, along with 3D-RRA identified substructures and Pfam domain annotations. The right panel shows the hotspot substructures in AlphaFold-predicted 3D DGKZ structure. The GOF-associated substructure is shown in blue and encircled with a dashed line. **(e)** Bar charts comparing the proportion of substructure harboring pathogenic mutations between significant and non-significant groups determined by 3D-RRA, in DDR proteins (upper) and T-cell regulatory proteins (bottom). P-values were calculated using Fisher’s exact test. **(f)** Heatmap showing three phenotypic clusters (LOF, blue; GOF, red; SOF, green) with functional mutations in PNKP. See legend of Fig. 2a for data description. The bottom color bar indicates mutation types (nonsense/splicing in purple, missense in gray). **(g)** AlphaFold-predicted 3D structure of PNKP, where residues harboring SOF-associated mutations are colored in green, and the corresponding substructure is highlighted with a dashed circle and a close-up view. In this substructure, three residues (R35, C46, Q50) carrying unscreened pathogenic mutation in ClinVar are highlighted in magentas.

Applying 3D-RRA to the DDR and T-cell regulator datasets, we identified 42 and 490 hotspot substructures (FDR<0.05), respectively. In both datasets, over 70% of these hotspot substructures overlap with annotated Pfam domains, facilitating the assignment of molecular functions. Consistent with the functional roles of DDR proteins and T-cell regulators, substructures in DDR proteins are enriched for enzymatic or DNA/RNA binding functions, whereas those in T-cell regulators primarily mediate protein-protein interactions (Fig. 3b, Supplementary Table S3). Notably, ProTiler-Mut also reveals novel functional substructures. As shown in Figures 3c and 3d, we identified four LOF-associated (Cluster 1, red) and one GOF-associated (Cluster 2, blue) hotspot substructures in DGKZ, a metabolic kinase negatively regulating T cell activation^37^. Four LOF-associated substructures align precisely with annotated DGKZ domains, where mutations predominantly impair cytokine secretion and surface marker expression^38^. Interestingly, a previously unannotated substructure spanning AA650-740 is enriched in mutations that exhibit a GOF phenotype, suggesting a distinct functional role in T cell regulation that warrants further investigation (Fig. 3d). ProTiler-Mut also uncovered an unannotated substructure in AA170-210 of the TRAIP ubiquitin ligase, where mutations specifically confer resistance to the TOP1 inhibitor camptothecin (CPT), recapitulating our recent findings^15^ (Fig. S3). Together, these results highlight the utility of ProTiler-Mut in not only corroborating known functional domains but also uncovering novel, potentially critical substructures that merit further exploration.

Since hotspot substructures are identified based on protein 3D structures, they can encompass distal amino acids that are brought into proximity through protein folding, defining functional units beyond the conventional annotation of linear protein domains. Among the hotspot substructures identified in the DDR and T-cell regulator datasets, 18 (41.9%) and 286 (58.3%) involve functional mutations at amino acids that are more than 100 residues apart in the primary sequence, respectively (Fig. S4a, S4b). For instance, we identified a hotspot substructure in XRCC3, a key protein in the homologous recombination pathway, that includes spatially proximal but sequence-distal residues E104, P282, and W288. Mutations of these residues sensitize cells to multiple DNA-damaging agents, suggesting an interplay between these residues in maintaining the activity of XRCC3 (Fig. S4c,4d).

In CRISPR-based mutagenesis screens, the requirement for a PAM sequence and the choice of base editors restricts the coverage of targetable sites. Additionally, many target sequences are inefficiently editable, limiting the sensitivity of these screens. The identification of hotspot substructures offers a way to connect phenotypic patterns in the screens to pathogenic mutations that are within these substructures but were not directly screened. Indeed, pathogenic mutations are highly enriched in hotspot substructures— approximately twice as much as in other regions deemed insignificant by statistical analysis of 3D-RRA (Fig. 3e). To facilitate future investigations, we mapped 55 and 341 pathogenic mutations in DDR proteins and T-cell regulators to the hotspot substructures identified in this study (Supplementary Table S4). For instance, Figure 3f shows a hotspot substructure in the DNA repair enzyme PNKP, located in the phosphorylated substrate-binding region of its FHA domain^39^, where mutations exhibit an SOF phenotype and specifically sensitize cells to the PARP inhibitor Olaparib (OLAP). Interestingly, three unscreened pathogenic mutations—R35G, C46G, and Q50E—implicated in neural disorders^40, 41^ are located in this substructure. This observation suggests that these mutations may contribute to disease development via an SOF mechanism. In support of this hypothesis, a previous study indicated that Q50E has minimal impact on its phosphatase and kinase activity. However, Q50E disrupts the nuclear localization of PNKP, leading to accumulation of single strand breaks in nuclear DNA but not in mitochondrial DNA^42^.

Collectively, our results demonstrate that identifying hotspot substructures enhances the interpretability of tiling mutagenesis screen data in three ways. First, it maps both known and novel protein domains to specific phenotypic patterns. Second, it uncovers the interplay between spatially proximal but sequence-distal residues that collaboratively influence protein activity. Third, it facilitates functional inference for pathogenic mutations within hotspot substructures, including those not directly screened or inefficiently inserted into the genome due to low sgRNA editing efficiency.

### ProTiler-Mut identifies protein-protein interactions disrupted by functional mutations

Proteins execute their functions through interactions with partners, forming intricate networks that regulate diverse cellular processes. Advances in AI-driven 3D structure prediction (e.g., AlphaFold-Multimer) and the expanding repository of experimentally solved structures have greatly enriched our understanding of protein-protein interaction (PPI) interfaces^43–45^. We reasoned that mapping functionally characterized mutations onto PPIs could provide insights into the mechanism underlying their impact on protein activities. To this end, ProTiler-Mut leverages functional mutations identified in mutagenesis screens as ‘anchors’ to systematically scan experimentally solved or AI-predicted complex structures, thereby identifying protein partners whose interactions may be disrupted. This strategy aids in prioritizing key mutations and candidate interaction partners for further experimental validation (Fig. 4a).

**Figure 4.**
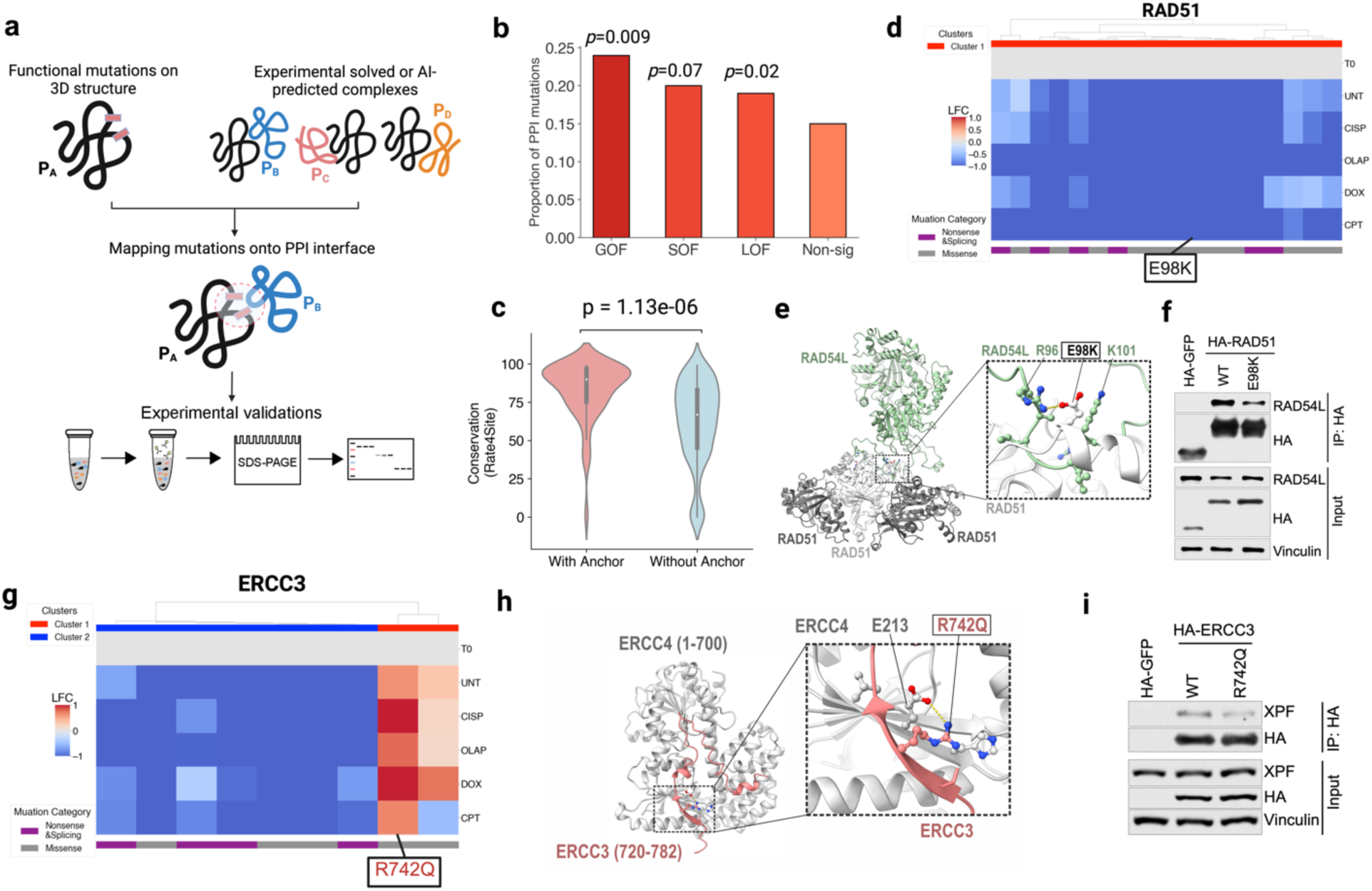
ProTiler-Mut identifies mutation-disrupted protein–protein interactions (PPI). **(a)** Schematic illustration of the ProTiler-Mut workflow for identifying mutational-disrupted PPIs. **(b)** Bar chart comparing the proportion of putative PPI-disrupting mutations in LOF, SOF and GOF functional mutation categories versus phenotypically non-significant (Non-sig) mutations. Analysis was based on BE tiling screens targeting 86 DDR proteins^15^. The p-values were determined using Fisher’s exact test. **(c)** Violin plot comparing the conservation scores (Rate4Site) for residues in predicted PPI interfaces with or without anchoring functional mutations. P-value was calculated using the Manny-Whitney U-test. **(d)** Heatmap showing the cluster of LOF mutations in RAD51, where E98K was predicted to disrupt the RAD51–RAD54L interaction and is highlighted with a rectangular box. **(e)** 3D structural model of the RAD51–RAD54L complex, with a close-up view of the interaction interface. E98K and its surrounding residues are shown in stick representation. **(f)** Co-IP assays showing the effect of E98K on RAD51-RAD54L interaction in HEK293T cells. **(g)** Heatmap depicting two clusters of functional mutations (LOF, blue; GOF, red) in ERCC3. The GOF mutation R742Q was predicted to disrupt ERCC3-ERCC4 interaction and is highlighted with a rectangular box. **(h)** 3D structural model of ERCC3 (720-782) in complex with ERCC4 (1-700), with a close-up view of the interaction interface. R742 residue and surrounding residues are shown in stick representation. **(i)** Co-IP assays showing the effect of R742Q on the interaction of ERCC3 and ERCC4/XPF in HEK293T cells.

As a proof-of-concept application, we mapped 522 functional mutations from the DDR dataset to 40,456 complex structures in the Predictome database^46^. Of these, 119 mutations (22.8%) were found within at least one predicted PPI interface with high confidence (average PAE < 5, pDOCKQ > 0.5). Among them, 74 (62.2%) were LOF, 19 (16.0%) were GOF, and 17 (14.3%) were SOF (Supplementary Table S5), suggesting that while most PPI-disrupting mutations impair protein activity, some may interfere with PPIs that negatively regulate protein function. Despite their differing functional consequences, all three mutation categories (LOF, GOF, and SOF) showed enrichment in PPI interfaces compared to mutations that did not exhibit strong phenotypes in the screens (Fig. 4b). Furthermore, PPI interfaces harboring functional mutations are more evolutionarily conserved than those without mutations, emphasizing their functional significance (Fig. 4c). This finding supports the strategy of using functional mutations as “anchors” to probe predicted PPIs.

Several of these PPI-associated functional mutations have been reported to disrupt critical PPIs. For instance, the D28N mutation in RIF1, which exhibits an SOF phenotype by specifically sensitizing cells to DOX treatment, was found at the interface between RIF1 and SHLD3 (Fig. S5a). A previous study demonstrated that this mutation abolishes the RIF1-SHLD3 interaction, preventing the recruitment of shieldin to DNA damage sites and thereby impairing DNA repair^47^. Another example involves BARD1, where ProTiler-Mut identified three PPI-associated LOF mutations—D712N, V713I, and D729N (Fig. S5b). Among them, D712 and V713 are key residues for the formation of the BARD1– histone H2B complex, while D729 is essential for BARD1’s interaction with ubiquitin, as supported by recent mechanistic studies^48^.

To further evaluate our method, we selected four PPI-associated functional mutations for experimental validation using co-immunoprecipitation (co-IP) (see Methods). Three of these mutations reduced binding affinity with their respective partners (Figs. 4f, 4i, S6), while one did not (Fig. S7). None of these mutations affected protein stability. Two of the three validated mutations—E98K in RAD51 and P10S in RAD51D—were classified as LOF (Figs. 4d, S6a) and impaired the RAD51-RAD54L and RAD51D-HELQ interactions, respectively (Figs. 4e, 4f, S6b, S6c). Both mutations reduced cell fitness upon treatment with multiple DNA-damaging agents, suggesting the critical role of these interactions in maintaining DNA repair fidelity. Intriguingly, the third mutation, R742Q in XPB/ERCC3, weakened the ERCC3-ERCC4 interaction, while exhibiting a GOF phenotype in the BE screen (Fig. 4g). While the ERCC3-ERCC4/XPF interaction is essential for nucleotide excision repair (NER)^49^, ERCC3 also functions in transcription initiation as a core component of the TFIIH complex, which regulates cell fitness independently of ERCC4^50,51^. This dual functionality may explain the observed GOF phenotype of R742Q. Although further investigation is needed to clarify how R742Q-mediated PPI disruption leads to a GOF effect, this case highlights the intricate balance of PPI dynamics in multi-functional proteins and the potential for compensatory mechanisms following interaction loss.

### Streamlined analysis with ProTiler-Mut identified pathogenic GOF mutations in MAPK1 that disrupt MAPK1-RSK1 interaction

To facilitate systematic analysis of tiling mutagenesis screen data, ProTiler-Mut was developed as a streamlined framework integrating multiple computational modules including pre-processing, functional mutation calling, mutation classification, 3D-RRA analysis, and PPI-mutation mapping (Fig. S8). ProTiler-Mut generates comprehensive annotations for functional mutations, hotspot substructures and PPI-mutation associations. It also provides 1D and 3D visualizations of proteins and functional mutations. Here, we illustrate how this streamlined analysis uncovered novel insights into pathogenic mutations in MAPK1/ERK2, a key signaling protein involved in regulating various cellular processes.

MAPK1 is activated through a phosphorylation cascade initiated by Ras-Raf-MEK-ERK signaling. Dysregulation of this pathway is associated with cancer, inflammatory disease, and neurodegenerative disorders^52–54^. In primary T cells, MAPK1 plays a key role for T cell activation and checkpoint regulation including PD-1 expression^55^. Thus, MAPK1 is one of the regulators targeted in the T cell BE dataset. ProTiler-Mut analysis identified two classes of MAPK1 mutations with LOF (Cluster 1) and GOF (Cluster 2) phenotypes, respectively (Fig. 5a). The GOF mutations predominantly reside in Y131 and Y316, with recurrent cancer mutations enriched in their surrounding amino acids (Fig. 5b). Although 185 AAs apart in the linear sequence, the two residues are spatially proximal within the MAPK1 structure, forming a hotspot substructure (Fig. 5c). This substructure also harbors two unscreened pathogenic mutations D318G and E322K. Fluorescence-activated cell sorting (FACS) analysis confirmed that D318G, E322K, Y131C, and Y316H all elevate PD-1 expression in HEK293T/PD-1 and Jurkat T cells (Figs. 5d and S9a), demonstrating consistent phenotypic effects across different cell types. The GOF phenotype was further supported by western blot analysis that showed increased MAPK1 phosphorylation in these mutants (Figs. 5e, S9b).

**Figure 5.**
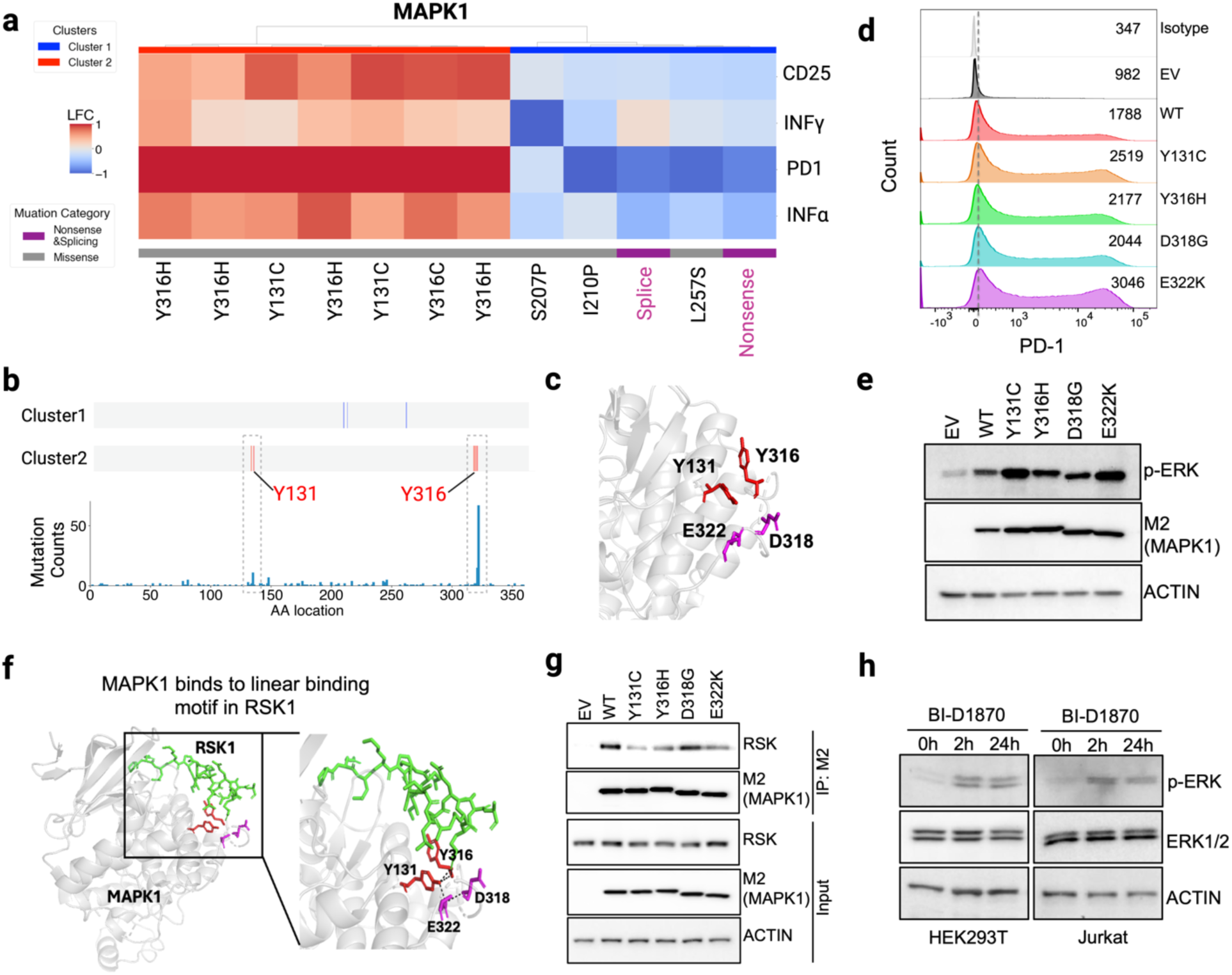
ProTiler-Mut identifies pathogenic GOF mutations that lead to MAPK1/ERK2 activation and elevated PD-1 expression. **(a)** A heatmap showing two clusters (LOF, blue; GOF, red) of functional mutations in MAPK1. See legend of Fig. 3c and main text for details of data source and color bar representation. **(b)** Distribution of mutations in clusters identified in (a) along the MAPK1 protein sequence, aligned with recurrent mutation frequencies (bottom track) retrieved from the COSMIC cancer database. **(c)** AlphaFold-predicted 3D structure of GOF-associated hotspot substructure in MAPK1, where residues harboring GOF mutations and two unscreened pathogenic mutations (E322K, D318G) are highlighted in red and pink, respectively. (**d-e**) Analysis of PD-1 surface expression and MAPK1 phosphorylation. HEK293T cells were co-transfected with PD-1 and either empty vector (EV), wild-type (WT) or mutant MAPK1. **(d)** PD-1 surface levels were assessed by FACS, and representative histograms from three independent experiments with mean fluorescence intensity (MFI) values are shown **(e)**. MAPK1 phosphorylation and expression of transfected constructs were evaluated by western blot. **(f)** Structural model of MAPK1 (gray) in complex with a linear motif in RSK1 (green, PDB: 3TEI), with a close-up view of the interaction interface. **(g)** Co-IP assay showing the effects of the MAPK1 mutants on MAPK1-RSK1 interaction in HEK293T cells. **(h)** HEK293T or Jurkat cells were treated with 10 μM RSK inhibitor BI-D1870 for the indicated time points. Total cell lysates were collected and analyzed by western blot.

Given that the hotspot substructure is located on the surface of MAPK1, we hypothesized that these mutations regulate MAPK1 activity by disrupting its interaction with binding partners. Using the hotspot substructure as an anchor, we scanned MAPK1-containing complex structures in the Protein Data Bank (PDB) and identified three partners—RSK1, MKP3, and MNK1—that bind to this region (Fig. 5f, Fig. S10a, 10b,). Among them, RSK1 has been reported by two independent studies to negatively regulate MAPK1 activity^56, 57^.

We therefore tested whether the mutations disrupt MAPK1–RSK1 interaction. Co-IP experiments confirmed that binding affinity between MAPK1 and RSK1 were impaired in the mutants (Fig. 5g). Furthermore, treatment with the RSK1 inhibitor BI-D1870 led to increased MAPK1 phosphorylation, supporting the role of RSK1 as a negative regulator of MAPK1 activity (Fig. 5h). Collectively, the streamlined ProTiler-Mut analysis, along with experimental validation, revealed that pathogenic mutations D318G and E332K, together with other mutations in the identified hotspot substructure, enhance MAPK1 activity and PD-1 expression by disrupting the MAPK1-RSK1 interaction, which otherwise restrains MAPK1 phosphorylation.

### Applicability of ProTiler-Mut to PE, DMS, and HDR-mediated SGE data

Prime editing (PE) offers unique flexibility in introducing targeted insertions, deletions, and substitutions^58, 59^. To assess its applicability to PE-based screens, we applied ProTiler-Mut to a recent PE tiling mutagenesis screen targeting *EGFR* in PC9 non-small cell lung cancer (NSCLC) cells treated with the tyrosine kinase inhibitors afatinib or osimertinib^26^. As shown in Fig. 6a, clustering analysis identified two key resistance groups: a common resistance mutation cluster (Cluster 1, red) and an afatinib-specific resistance cluster (Cluster 2, blue). The common resistance mutations localize to two hotspot substructures—AA715-730 near the drug-binding pocket and AA915-935, a potential distal allosteric site identified in a previous study^21^ (Fig. 6b). In contrast, Cluster 2 mutations, specific to afatinib resistance, are confined to the afatinib-binding pocket (Fig. 6c), which includes the well-established drug resistance mutation T790M^60^. Consistent with this pattern, osimertinib was developed to overcome T790M-mediated resistance^61^, explaining why mutations in Cluster 2 have minimal impact on cellular sensitivity to osimertinib.

**Figure 6.**
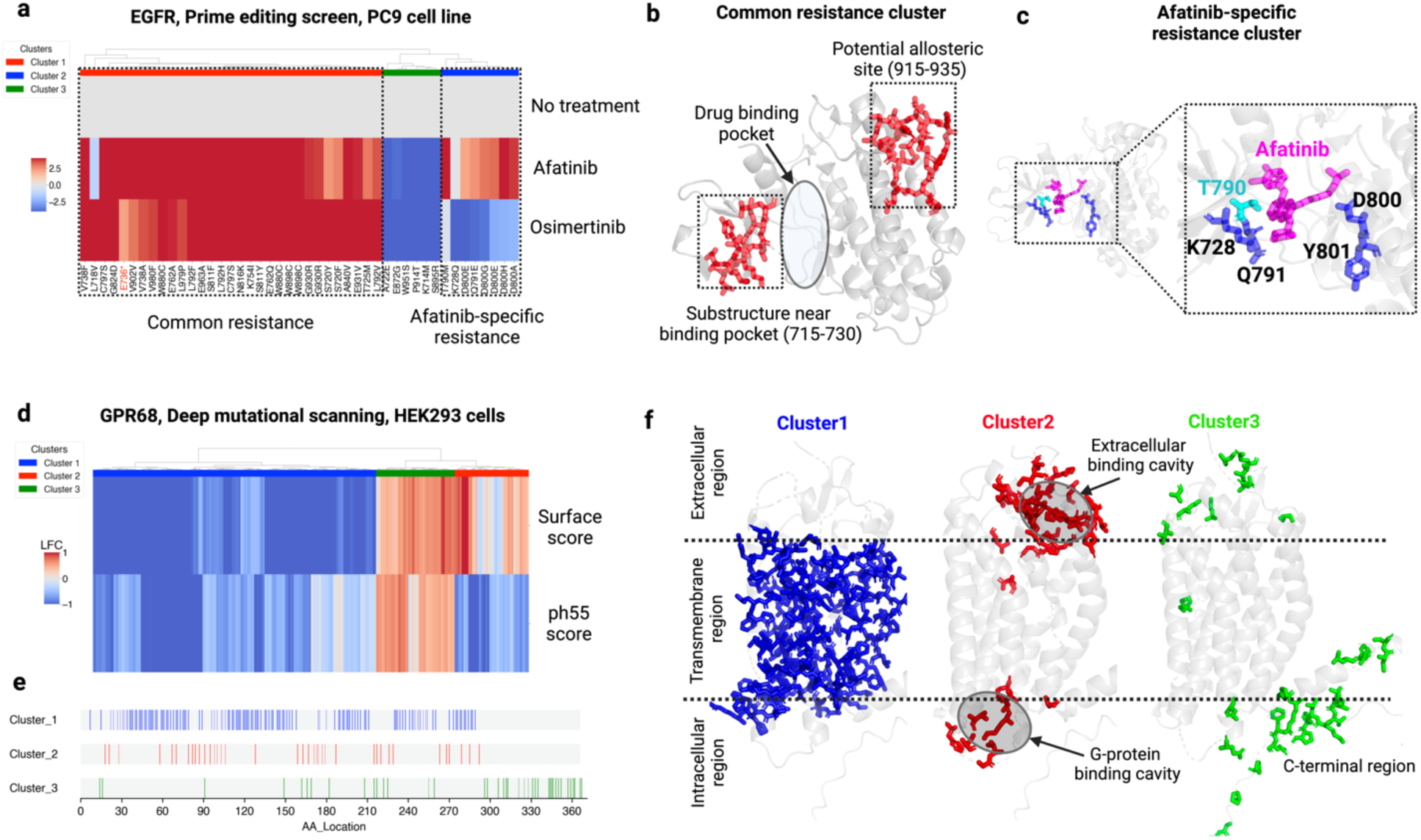
ProTiler-Mut is applicable to PE, DMS tiling mutagenesis screens. **(a)** A heatmap showing three phenotypic clusters (common resistance, red; afatinib-specific resistance, blue; no resistance, green) of functional mutations in EGFR that mediate PC9 cell response to afatinib or osimertinib. Data retrieved from published PE tiling screen^26^ **(b)** A structural model of EGFR (PDB: 4G5J) highlighted with two hotspot substructures enriched with mutations leading to common resistance. **(c)** A structural view of mutations leading to afatinib-specific resistance (T790M in cyan and others in blue) and the afatinib molecule (pink). The close-up view shows the hotspot substructure region. **(d)** Heatmap showing three phenotypic clusters of residues in GPR68 (see main text for details). Data was retrieved from a published DMS screen targeting GPR68^13^ . **(e)** Distribution of amino acids in different clusters identified in (d) along the protein sequence of GPR68. **(f)** AlphaFold-predicted structure of GPR68 overlaid with the residues in each phenotypic cluster in (d). Two dashed lines denote the membrane boundaries, separating the extracellular, transmembrane, and intracellular regions. The extracellular binding cavity and the G-protein binding cavity in cluster 2 are highlighted with gray circles.

We also applied ProTiler-Mut to a recent deep mutational scanning (DMS) dataset focusing on variants of GPR68^13^, a proton-sensing G protein-coupled receptor crucial for detecting extracellular pH changes. The DMS screen assigned two phenotypic scores to each variant: a surface score that reflects the impact of the variant on GPR68 cell surface expression, and a ph55 score that assesses GPR68 activity under acidic extracellular conditions. After aggregating the variant scores by residue, clustering analysis identified three distinct groups: Cluster 1 (blue), associated with low surface expression and low pH55 scores; Cluster 2 (red), associated with high surface expression but low pH55 scores; and Cluster 3 (green), associated with high scores for both (Fig. 6d). While residues in each cluster were distributed across the linear amino acid sequence (Fig. 6e), mapping these residues to the GPR68 structure revealed distinct localization patterns (Fig. 6f). Residues in Cluster 1 are predominantly located within the transmembrane helices, whereas those in Cluster 2 and 3 are found in the extracellular and intracellular regions. Notably, residues in Cluster 2 define two hotspot substructures, corresponding to the extracellular binding cavity and the G-protein binding cavity, which are implicated in acidic sensing and signal transduction but do not affect cell surface expression^13^.

Finally, we applied ProTiler-Mut to analyze two independent HDR-mediated saturation genome editing (SGE) datasets targeting RAD51C^19^ and VHL^18^, respectively. Since these SGE screens were designed with a single condition (cell viability), only GOF and LOF mutations were identified (Fig. S11a and S11b). Despite this limitation, ProTiler-Mut successfully identified a hotspot substructure in RAD51C, where GOF mutations enhance the binding between RAD51C and RAD51D by increasing hydrophobic interactions at the PPI interface^19^ (Fig. S11a). ProTiler-Mut also highlighted GOF mutations in the N-terminal tail of VHL, which mediate its interaction with the tumor suppressor p14ARF^62^ (Fig. S11b). Collectively, these analyses demonstrated the versatility of ProTiler-Mut, showing its compatibility with various scoring matrices and its broad applicability across diverse tiling mutagenesis screening platforms.

## DISCUSSION

With advances in CRISPR technology, high-throughput mutagenesis screens have become powerful tools for mapping protein mutations to phenotypic outcomes at single– amino acid resolution. However, these screens face challenges related to limited coverage, sensitivity, and statistical power. ProTiler-Mut addresses these limitations by leveraging a tiling mutagenesis design that systematically perturbs amino acid contexts across entire proteins. By integrating multiple mutations that exhibit similar phenotypic patterns and cluster in spatially proximal residues, ProTiler-Mut enhances analytical fidelity and enables the inference of functions for unscreened or low-confidence mutations.

When applied to screens with multiple conditions, ProTiler-Mut analysis identified SOF mutations, expanding beyond the conventional binary classification of LOF and GOF mutations used in previous mutagenesis screen analysis. Notably, SOF mutations in DDR proteins and T cell regulators exhibited a higher enrichment of pathogenic mutations compared to LOF and GOF variants. Indeed, several pathogenic SOF mutations have been characterized in previous studies. For instance, pathogenic SOF mutations in TP53 selectively impair DNA damage-dependent apoptosis without affecting cell-cycle arrest, specifically contributing to cancer progression^63^. Similarly, pathogenic SOF mutations in STAT3 have been associated with hyper-IgE syndrome by selectively disrupting cytokine-induced immune signaling pathways^64^. These examples highlight the significance of SOF mutations in multi-functional proteins, where a partial loss of activity, rather than complete inactivation, can drive disease in an environment-dependent manner. Importantly, detecting SOF mutations is uniquely achievable through mutagenesis screens, as traditional knockout or knockdown approaches lack the resolution to capture these nuanced functional shifts.

ProTiler-Mut also proves useful for predicting PPIs disrupted by functional mutations. This capability is powered by recent advancements in AI-predicted complex structures. To optimize computational efficiency, the current version of ProTiler-Mut relies on previously predicted or experimentally solved structures and is focused on spatial proximity between mutated residues and their interaction sites with partner proteins. Moving forward, ProTiler-mut will integrate state-of-the-art structural prediction and docking algorithms^45, 65^ for further improvement of accuracy. This future iteration will also extend its capabilities to include protein interactions with DNA, RNA, ions, small molecules, and modified residues.

Recently, high-throughput mutagenesis screening platforms have seen significant optimization and expansion. For instance, sensor-based reporter systems have enhanced the interpretability of BE and PE screens by capturing precise editing outcomes^25, 66^. Moreover, single-cell mutagenesis screens are emerging as a promising technology^67–69^. ProTiler-Mut, designed to be compatible with various mutagenesis platforms, offers the flexibility to seamlessly integrate with these evolving technologies. This adaptability ensures a more comprehensive analysis of mutation-driven alterations in protein functionality in the future.

## METHOD

### ProTiler-Mut Pipeline

ProTiler-Mut is a comprehensive computational framework designed to analyze tiling mutagenesis screening data, focusing on the identification, visualization, and functional characterization of significant missense mutations in protein-coding regions. The pipeline integrates multiple computational modules including pre-processing, functional mutation calling, mutation classification, 3D-RRA analysis, and PPI-mutation mapping, providing comprehensive structural, functional as well as mechanistic insights into the identified mutations in the screen data.

### Inputs

The primary input of ProTiler-Mut is a meticulously curated table generated from tiling mutagenesis screens which include the following key information: i) the exact location within the protein sequence where the mutation is introduced; ii) the functional nature of the mutation (*e.g.,* missense, nonsense, splicing), and iii) the specific amino acid change (*e.g.* R106K). The table should also contain experimentally measured phenotypic effects associated with each mutation (*e.g.,* raw read counts, log fold changes, z-scores, or other phenotypic metrics) in at least one experimental condition. In BE or PE screens, each guide RNA corresponds to an entry (row) in the table such that two or more guide RNAs introducing the same mutation are not aggregated and the experimental measurements in the screen are assigned separately. Additional metadata (*e.g.,* experimental conditions, sample identifiers, or assay parameters) can also be included to offer further context.

### Preprocessing & functional mutation calling

If raw read counts are provided, ProTiler-Mut normalizes the data across multiple conditions to correct for disparities in sequencing depth and batch effects. This normalization assumes most mutations do not produce a detectable phenotype, so each mutation’s read count (*R*_*i*_) is divided by the median read count of all mutations *median*(*R*) in that condition:

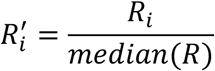

where 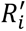 is the normalized read count for the mutation *i* . Note that ProTiler-Mut is designed to work with pre-processed tabular data; users are required to convert raw FASTQ files into the appropriate table format using other software (*e.g.* MAGeCK) before using the pipeline.

Next, ProTiler-Mut will calculate the Log Fold Change (LFC) as phenotypic score *x*_*i*_by comparing 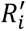 to the normalized read counts 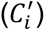 in the control condition (e.g. pre-selection condition, plasmid control, etc.)

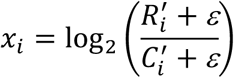

Where ε is a small positive value to avoid large variation due to low read counts, and the default value was set to be 0.05.

ProTiler-Mut then identifies significant mutations by benchmarking their phenotypic score against those observed in negative control mutations. These controls can include sgRNAs targeting neutral loci (*e.g*., AAVS), non-targeting sgRNAs, or synonymous mutations that do not alter the amino acid sequence. Specifically, ProTiler-Mut calculates a Z-score for each mutation using the formula:

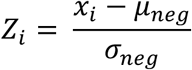

where 𝜇_neg_, 𝜎_neg_ denote the mean and standard deviation of the negative control distribution, respectively.

For those high-density screens (e.g. Deep mutational scan), where each amino acid has all the possible mutations introduced in the screen, ProTiler-Mut averaged the phenotypic effects of all the mutations for a specific amino acid as *xA*_*i*_, followed by calculation of Z-score:

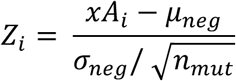

Where 𝑛_*mut*_ is the number of mutations introduced for the specific amino acid.

If multiple conditions (𝑐 = 1,2 …, 𝑛) are provided, the pipeline uses the maximum difference across all conditions to capture the strongest phenotypic effect. Concretely, the maximum difference (Δ_*max*,*i*_) for mutation *i* is calculated as:

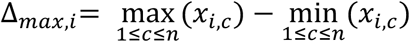

where *x*_*i,c*_ is the observed or normalized effect of mutation *i* under condition 𝑐 . Finally, ProTiler-Mut calculates a corresponding Z-score by comparing each Δ_*max*,*i*_value to those from negative control mutations, thereby reflecting the potential functional significance of each mutation under one or more conditions. Significant mutations are selected by evaluating the Z-scores calculated for each mutation. In essence, the Z-score quantifies how far the observed (or normalized) phenotypic effect of a mutation deviates from the mean effect of the negative controls, relative to the standard deviation of that control distribution. Typically, mutations with a Z-score greater than 2 or less than -2 are considered statistically significant, indicating that their effects are more than two standard deviations away from the baseline noise. This threshold, however, can be adjusted based on dataset-specific characteristics such as variability and noise levels, ensuring an optimal balance between sensitivity and specificity in identifying functionally relevant mutations.

### Clustering and Mutation categorization

Given the set of significant mutations and their corresponding phenotypic scores under different conditions, ProTiler-Mut next performs hierarchical clustering on the resulting matrix, where each row represents the scores for a specific treatment condition, and each column corresponds to one of the identified significant mutations. The clustering parameters—such as the linkage method and distance metric—are adjustable based on the nature of the dataset. For instance, when only one condition is available (as is common in viability screens), we use Euclidean distance with Ward’s linkage. In contrast, for datasets with multiple treatment conditions (*e.g.,* different drug treatments), we employ correlation-based distance with average linkage, allowing ProTiler-Mut to capture more nuanced relationships across multiple experimental settings.

For screens performed under a single condition, ProTiler-Mut employs a direct comparison strategy to classify mutations. For a given mutation *i*, let *x*_*i*_denote its observed phenotypic score under the single experimental condition. First, the average phenotypic score of the reference set—comprising known loss-of-function mutations (nonsense and splicing mutations)—is computed as:

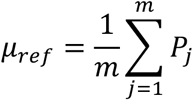

Where {𝑃_1_, 𝑃_2_, …, 𝑃_*m*_} represents the phenotypic scores of the reference mutations. The classification rule is then defined as follows:

- If *Sign*(*x*_*i*_) = *Sign*(𝜇_ref_) (*i. e*., *x*_*i*_ × 𝜇_ref_ > 0), the mutation *i* is classified as LOF.
- If *Sign*(*x*_*i*_) ≠ *Sign*(𝜇_ref_) (*i. e*., *x*_*i*_ × 𝜇_ref_ < 0), the mutation *i* is classified as GOF.

If the screen was performed under multiple condition, we compare the phenotypic effects of a specific mutation *i* across multiple conditions (𝑐 = 1,2 …, *n*) to those of a reference set of *m* nonsense and splicing mutations. Let:

- 𝑉_*i*_ be the score vector for mutation *i*, of length *n* (one entry per condition),
- 𝑆_ref_ = {𝑃_1_, 𝑃_2_, …, 𝑃_*m*_} be the set of score vectors for all nonsense or splicing mutations targeting specific protein, each also of length *n*.

We then calculate the average correlation between 𝑉_*i*_ each 𝑃_j_ in 𝑆_ref_ . Let 𝑐𝑜𝑟𝑟(𝑉_*i*_, 𝑃_j_) represent the Pearson’s correlation between 𝑉_*i*_ and 𝑃_j_ . We then define the average correlation for mutation *i* as:

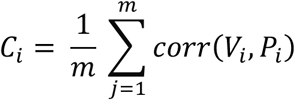

ProTiler-Mut leverages the average Pearson correlation coefficient, 𝐶_*i*_, to quantify how closely the phenotypic profile of mutation *i* resembles that of the reference set of nonsense and splicing mutations. This measure is then used to categorize mutation *i* as loss-of-function (LOF), separation-of-function (SOF), or gain-of-function (GOF) based on the global distribution of all 𝐶_*i*_values across the dataset. In practice, if 𝐶_*i*_< 0, indicating that the mutation’s effect is anti-correlated with the reference mutations, it is classified as GOF; if 𝐶_*i*_ >0.5, reflecting a high degree of correlation with the reference mutations, it is classified as LOF; and if 𝐶_*i*_ falls between these thresholds, the mutation is categorized as SOF, showing moderate correlation with the reference set. It is important to note that these threshold values serve as general guidelines rather than absolute cutoffs. Therefore, the functional classification of an individual mutation should be interpreted by integrating both the quantitative correlation metrics and the qualitative phenotypic patterns observed in detailed heatmap visualizations, ensuring a comprehensive assessment of its biological impact. Moreover, the precision of the correlation analysis is influenced by the number of conditions included in the screen—generally, incorporating more conditions yields a more robust and reliable correlation assessment.

### Amino acid-level annotations

ProTiler-Mut provides comprehensive amino acid-level annotations to elucidate the biological impact of functional mutations by integrating diverse external databases and predictive tools. Protein domain annotations were derived from the Pfam database^70^ (the current released version downloaded from the official FTP site https://ftp.ebi.ac.uk/pub/databases/Pfam/), which is used to map identified mutations onto known functional domains of the targeted protein. Post-translational modifications (PTMs) annotations data were downloaded from PhosphoSitePlus^71^ official website (https://www.phosphosite.org/staticDownloads). which is used to map identified mutations onto known phosphorylation, acetylation, methylation sites and other modification sites. Surface accessibility metrics, including residue depth and accessible surface area (ASA), are calculated using DSSP function in PDB module in Biopython^72^, providing quantitative insights into the solvent exposure of mutated residues. To assess protein stability, ProTiler-Mut leverages DDGun^33^ and RasP^32^, both of which are installed from their respective GitHub repositories (https://github.com/biofold/ddgun and https://github.com/dohlee/rasp-pytorch), to predict ΔΔG values and infer stability changes induced by mutations. Conservation scores are derived using Rate4Site^73^, which is run on multiple sequence alignments obtained from public databases, and by incorporating confidence metrics from AlphaFold predicted pLDDT scores. Additionally, pathogenicity annotations are integrated by mapping mutations to clinically relevant databases, including ClinVar^74^ and COSMIC^75^. ClinVar data was downloaded from https://ftp.ncbi.nlm.nih.gov/pub/clinvar/ and COSMIC data was downloaded from https://cancer.sanger.ac.uk/cosmic/download/cosmic. By automating the download, parsing, and integration of these resources, ProTiler-Mut systematically annotates each amino acid position, thereby providing detailed functional insights into the impact of functional mutations identified in the tiling mutagenesis screens.

### 3D-RRA module

ProTiler-Mut employs the 3D-RRA method, which integrates spatial mutation mapping with a Robust Rank Aggregation (RRA) algorithm to robustly identify functional substructures within a protein.

First, significant mutations from a specific cluster identified from the clustering module were mapped onto the protein’s three-dimensional structure as seed mutations. The seed mutations were further merged based on their spatial distance. Let 𝑀 = {*m*_1_, *m*_2_, …, *m*_*n*_} be the set of seed mutations and let *d*(*i*, *j*) denote the spatial distance between mutations *m*_*i*_ and *m*_j_. The merging was performed through following steps:

- Starting from the distance matrix 𝐷, where 𝐷[*i*, *j*] = *d*(*i*, *j*) *for all i* ≠ *j*;
- Identify the pair (*i*, *j*) with minimum distance while there exist a pair (*i*, *j*) such that *d*(*i*, *j*) < *t* (*where t is the threshold, default value is* 10), then merge *m*_*i*_ and *m*_j_ into a new cluster 𝐶 = {*m*_*i*_, *m*_j_}
- Update the distance between the new cluster 𝐶 and any other mutation *m*_𝑘_ using:

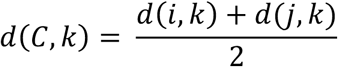

- Mark the individual mutation *m*_j_ as merged and not for further consideration Iterate the above process until no pair of mutation or clusters has a distance smaller than *t*, the final clusters consist of all merged mutation groups and any single mutations that were not merged.

Next, for each merged mutation group, ProTiler-Mut further include surrounding amino acid proximal to the seed mutation clusters to form a substructure. Let 𝖢 be the set of residue numbers corresponding to a seed mutation cluster and let ℛ be the set of all residues in the protein structure. For each residue *i* ∈ 𝖢 (provided *i* ∈ ℛ), let ℛ_*i*_ denote the residue in the structure and let 𝐴(ℛ_*i*_) be the set of its 3D atomic coordinates. For every residue R∈ ℛ, define the distance between ℛ _*i*_ and R as:

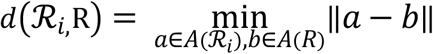

where ‖*a* − 𝑏‖ is the Euclidean distance between atoms *a* and 𝑏. A residue R is considered to be part of the substructure if:

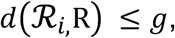

Where *g* is a predefined distance threshold, the default value is 5.

The overall substructure 𝑆 is then given by the union over all seed residues in 𝖢:

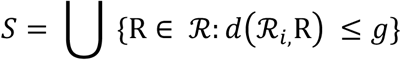

ProTiler-Mut then employed Robust Rank Aggregation (RRA) method to test if a given substructure is statistically significant. First, we ranked all the 𝑀 signals from the entire dataset based on their Z-scores, let *R* = (𝑟_1_, 𝑟_2_, …, 𝑟_*n*_) denote the vector of ranks of the n signals contained within that substructure. These ranks are normalized into percentiles 𝑈 = (𝑢_1_, 𝑢_2_, …, 𝑢_*n*_), where 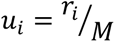. Under the null hypothesis, the percentiles are uniformly distributed between 0 and 1, and the 𝑘 -th smallest value among 𝑢_1_, 𝑢_2_, …, 𝑢_*n*_ follows a beta distribution 𝐵(𝑘, *n* + 1 − 𝑘). The RRA method then computes a *p*-value for each order statistic and defines the significance score as ρ = min (p_1_, p_2_, …, p_n_). To assign a final p-value to each substructure, we perform a permutation test in which signals are randomly assigned to specific substructure (maintaining the same number of signals) over 100 × N permutations (with N being the number of signals in whole dataset) Additionally, a false discovery rate (FDR) is estimated by ranking the permutation statistics and determining the relative position of the observed statistic, thereby offering an indication of the expected proportion of false positives among the findings. This integrated 3D-RRA method thus leverages both spatial context and rigorous statistical evaluation to efficiently identify functional protein substructures.

### PPI mapping module

To link the identified functional mutations or substructures to potential protein-protein interactions, ProTiler-Mut maps the functional mutation(s) to the 3D structure of PPI interface involving the target protein to identify potential PPI-associated mutations.

Let *m* be a functional mutation in target protein 𝑇, and denote the residue harboring this mutation as *R*_𝑇_ . Let 𝐴(*R*_𝑇_) be the set of 3D atomic coordinates of *R*_𝑇_ . For a partner protein 𝑃 composed of *n* residues {𝑃_1_, 𝑃_2_, …, 𝑃_*n*_}, let 𝐴(𝑃_*i*_) represent the set of 3D atomic coordinates for residue 𝑃_*i*_. The distance between *R*_𝑇_ and 𝑃_*i*_is defined as:

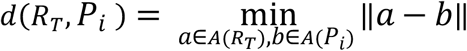

where ‖*a* − 𝑏‖ is the Euclidean distance between atoms *a* and 𝑏. The residue 𝑃_*i*_ is considered to interact with *R*_𝑇_ if:

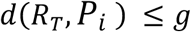

Where *g* is a predefined distance threshold, the default value is 5.

By iterating all the 𝑃_*i*_ in the partner protein, we will get a set of interacting residues as 𝐼 = {𝐼_1_, 𝐼_2_, …, 𝐼_𝑘_}, if 𝑘 > 0, *m* is defined as a PPI-associated mutation at the interaction interface between target protein 𝑇 and partner protein 𝑃. For a substructure containing multiple mutations, if at least one mutation is classified as PPI-associated, the entire substructure is assigned to the specific PPI interface.

The source of protein complex structures is flexible; users may input experimentally resolved structures from the PDB database or high-quality AlphaFold-predicted complexes (meeting metrics such as ipTM, average ipTM, PAE scoreS, etc.). By systematically scanning all user-provided protein complexes, ProTiler-Mut outputs all potential mutation-disrupted PPIs and prioritizes key residues and candidate interaction partners for further experimental validation.

In this study, 3D structures of protein complexes were obtained from the PDB database and from AlphaFold-based predictions available in the Predictome database^46^. Variants located at conserved, surface-exposed positions were examined. Four functional mutations associated with protein–protein interactions (PPIs) were selected for experimental validation based on their AlphaFold confidence scores, their presence within predicted interfaces in Predictome database, and the absence of the corresponding complex from existing entries in the PDB.

### Cell Culture and treatment

HEK293T cells were cultured in Dulbecco’s modified Eagle’s medium (Thermo Fisher, Cat. No. 11965092) supplemented with 10% fetal bovine serum (FBS) (Thermo Fisher, Cat. No. 10500056) while Jurkat cells were cultured in RPMI1640 medium (Thermo Fisher, Cat. No. 11875093) supplemented with 10% FBS. Cells were incubated in humidified incubators at 37oC and 5% CO2. Plasmid transfection was carried out using Lipofectamine™ 2000 (Thermo Fisher Scientific, Cat. No. 11668019) for HEK293T cells, or the SE Cell Line 4D-Nucleofector™ X Kit (Lonza, Cat. No. V4XC-1012) for Jurkat cells, following the respective manufacturer’s instructions. The RSK inhibitor BI-D1870 was obtained from MedChemExpress (Cat. No. HY-10510).

### Plasmids

RAD51 (NM_002875.5) and RAD51D (NM_002875.4) were amplified with flanking attB1/attB2 site sequences from HEK293T cDNA using the following primers: RAD51 (5’-ggggacaactttgtacaaaaaagttggcaccATGGCAATGCAGATGCAGCTTGA-3’ and 5’-ggggacaactttgtacaagaaagttgggCAGTCTTTGGCATCTCCCACTC-3’); RAD51D (5’-ggggacaactttgtacaaaaaagttggcaccATGGGCGTGCTCAGGGTC-3’ and 5’-ggggacaactttgtacaagaaagttgggCATGTCTGATCACCCTGTAATGTGGC-3’). Their respective amplicons were cloned into the Gateway entry vector pDONR223 using BP Clonase II (Thermo Fisher, Cat. No. 11789020). The pDONR223-ERCC3 plasmid from the hORFeome V8.1 library (clone ID: 4854) is a gift from the laboratory of Stephen Elledge (Brigham and Women’s Hospital, Harvard Medical School). Site-directed mutagenesis was performed via inverse PCR to introduce the following mutations:RAD51 E98K using primers 5’-AAGGCGGTCAaaaATCATACAGA-3’ and 5’-TGGTGGAATTCAGTTGCAG-3’; RAD51D P10S using primers 5’-CGGACTGTGCtCTGGCCTTAC-3’ and 5’-ACCCTGAGCACGCCCATG-3’; ERCC3 R742Q using primers 5’-CAGGCATCTCaGCGCTTTGGC-3’ and 5’-GCTGGATCTGGAGCCAAATTC-3’. To generate expression constructs, the wildtype and mutant entry vectors were recombined into the Gateway destination vector pMSCV-FLAG-HA-DEST using LR Clonase II (Thermo Fisher, Cat. No. 11791020). Expression plasmids pcDNA3.1-PD-1-cHA (Clone ID: OHu26320) and pcDNA3.1-MAPK1-M2 (Clone ID: OHu25335) were purchased from Genscript. MAPK1 point mutations were introduced by site-directed mutagenesis using PCR with the following primer pairs: Y131C (5’-TATTTTCTCTgCCAGATCCTCAG-3’ and 5’-GCAGATATGGTCATTGCTG-3’), Y316H (5’-TCTGGAGCAGcATTACGACCCGAG-3’ and 5’-TATGGGTGGGCCAGAGCC-3’), D318G (5’-CAGTATTACGgCCCGAGTGAC-3’ and 5’-CTCCAGATATGGGTGGGC-3’), E322K (5’-CCCGAGTGACaAGCCCATCGC-3’ and 5’-TCGTAATACTGCTCCAGATATGGG-3’). All constructs were verified by Sanger sequencing.

### Co-immunoprecipitation and western blotting

HEK293T were transfected with different expression constructs, and total lysates were harvested three days later after transfection for co-immunoprecipitation and western blot as described previously^76^. Briefly, the cells were harvested, washed with ice-cold PBS, and total cell extracts were prepared in lysis buffer (10mM Tris/Cl pH 7.5; 150mM NaCl; 0.5mM EDTA; 0.5% NP-40) supplemented with protease inhibitor (Sigma-Aldrich, Cat. No. 4693159001) and phosphatase inhibitor (Thermo Fisher, Cat. No. 78420) cocktails. the lysates were collected by centrifuge and incubated with HA-agarose beads (Sigma-Aldrich, Cat. No. A2095) or M2-agarose beads (Sigma-Aldrich, Cat. No. A2220) for 2 hours at 4°C for immunoprecipitation. Then the beads were washed three times with lysis buffer and boiled in loading buffer to elute the bounded proteins. The cell extracts or immunoprecipitated samples were separated by SDS-PAGE and transferred onto PVDF membranes. Blots were blocked in PBS containing 5% non-fat dry milk, then incubated with the appropriate primary antibody in the blocking buffer overnight at 4°C. The blots were then washed with PBST (PBS with 0.05% tween 20) and probed with an HRP-labelled secondary antibody for 1h at room temperature. After washing three times with PBST, the membranes were incubated with ECL reagent for immunoblotting.

The antibodies used for immunoblotting were: RAD54 (Cell Signaling, Cat. No. 15016, 1:1000 dilution), HELQ (Cell Signaling, Cat. No. 19436, 1:500 dilution), XPF (Fortis Life Sciences, Cat. No. A301-315A, 1:1000 dilution), HA (Sigma-Aldrich, Cat No. H3663, 1:5000 dilution), p-ERK (Cell Signaling, Cat. No. 9101, 1:1000 dilution), ERK1/2 (Cell Signaling, Cat. No. 4695, 1:1000 dilution), RSK (Cell Signaling, Cat. No. 9355, 1:1000 dilution), M2 (Sigma-Aldrich, Cat. No. F3165, 1:3000 dilution), actin (Sigma-Aldrich, Cat. No. 1978, 1:3000 dilution), vinculin (Sigma-Aldrich, Cat No. V9131, 1:20,000 dilution), and GAPDH (Bethyl, Cat. No. A303-878A, 1:2000 dilution).

### Flow cytometry (FACS) analysis for cell surface PD-1

The single-cell suspensions were generated in PBS buffer with 1% FBS and 0.1 mM EDTA. Cells were incubated with PE-conjugated anti-human PD-1 antibody (BD Biosciences, Cat. No. 560798) for 30 min at 4 °C in the dark, then the cells were washed twice with PBS buffer containing 1% FBS and 0.1 mM EDTA. Flow cytometry data were acquired on Fortessa and analyzed with Flowjo 10.10.0 software.

## CODE AVAILABILITY

ProTiler-Mut is publicly available at https://github.com/MDhewei/ProTiler-Mut. All the analyses presented in this manuscript were produced with version 0.1.0 (initial submission)

## ACKNOWLEDGEMENT

We thank Drs. Yiwen Chen and Taiping Chen for critical discussion. Work in the Han Xu laboratory was supported by NIH R35GM137927 and John S Dunn Research Foundation Award. Work in the Mark Bedford laboratory was supported by NIH R35GM153387. Work in the Alberto Ciccia laboratory was supported by NIH R01CA197774. H.X. is a CPRIT Scholar in Cancer Research.

## AUTHOR CONTRIBUTIONS

H.X., A.C., R.G. and W.H. conceptualized the study. W.H. developed the software. R.G. and H.W. performed the computational analysis. J.W.H. performed the experimental validation on predicted mutation-disrupted PPIs. Y.W. performed the experimental validation on the MAPK1 study. S.B.H., G.L., A.V. and R.F. helped dataset curation and analysis. R.G., G.L. and S.W. helped software development. M.T.B. helped data interpretation. H.X. and A.C. supervised the project. All authors participated in writing the manuscript.

