## Supplementary Figures for "Mechanistic Insights into Proteomic Mutation-Phenotype Linkages from Tiling Mutagenesis Screens"

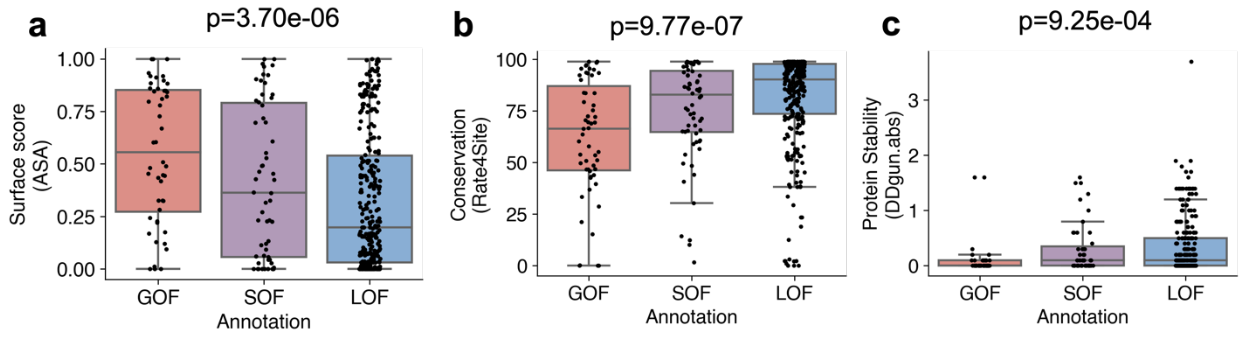


**Figure S1.** Box plots comparing amino acid features in addition to Fig.2g-2i: (a) Surface score (ASA: Accessible Surface Area); (b) evolutionary conservation scores (Rate4Site) and (c) free energy change (absolute DDgun score) among GOF, SOF, and LOF categories. P-values were calculated using the Kruskal–Wallis test, a non-parametric alternative to one-way ANOVA.

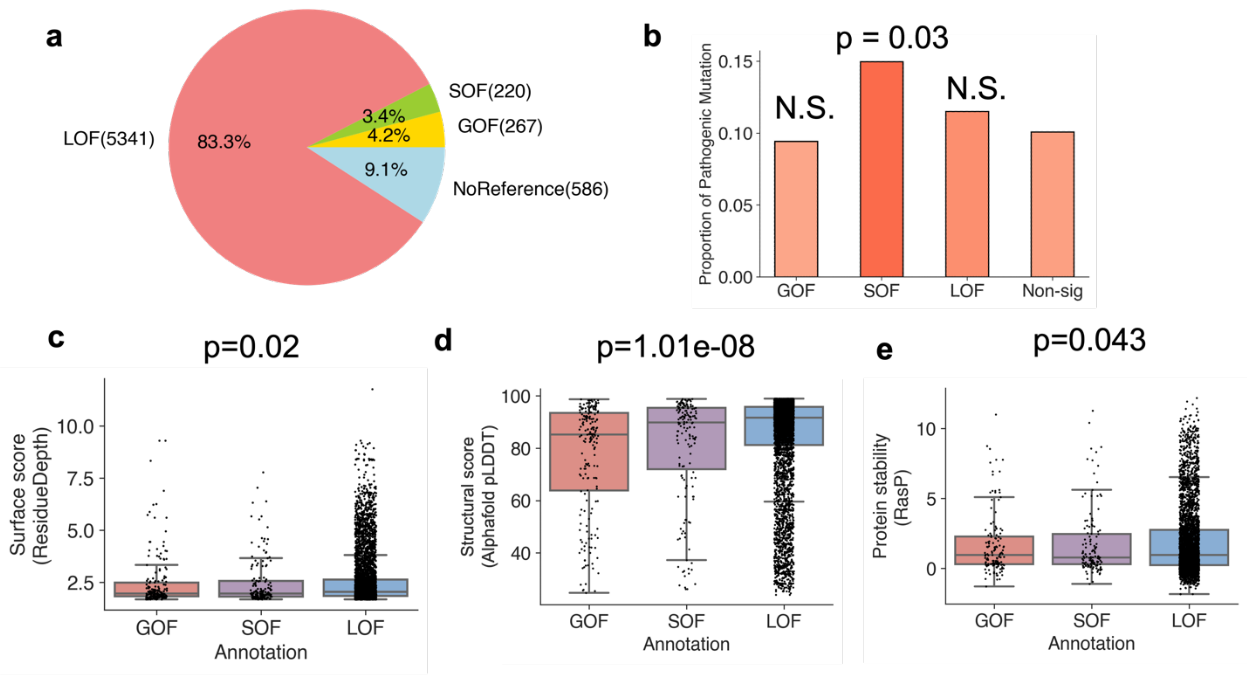


**Figure S2. Classification of functional mutations in T-cell dataset (a)** Pie chart showing the distribution of mutation categories in T-cell regulatory proteins. **(b)** Bar chart showing the proportion of pathogenic mutations in GOF, SOF, LOF categories, compared to mutations not associated with significant phenotype in the screen (Non-sig). P-values were calculated using Fisher's exact test (N.S.: not significant). **(c-e)** Box plots comparing (c) Surface score (the Residue Depth); (d) Structural scores (Alphafold score) and (e) free energy change (RasP score) among GOF, SOF, and LOF categories. P-values were calculated using the Kruskal–Wallis test, a non-parametric alternative to one-way ANOVA.

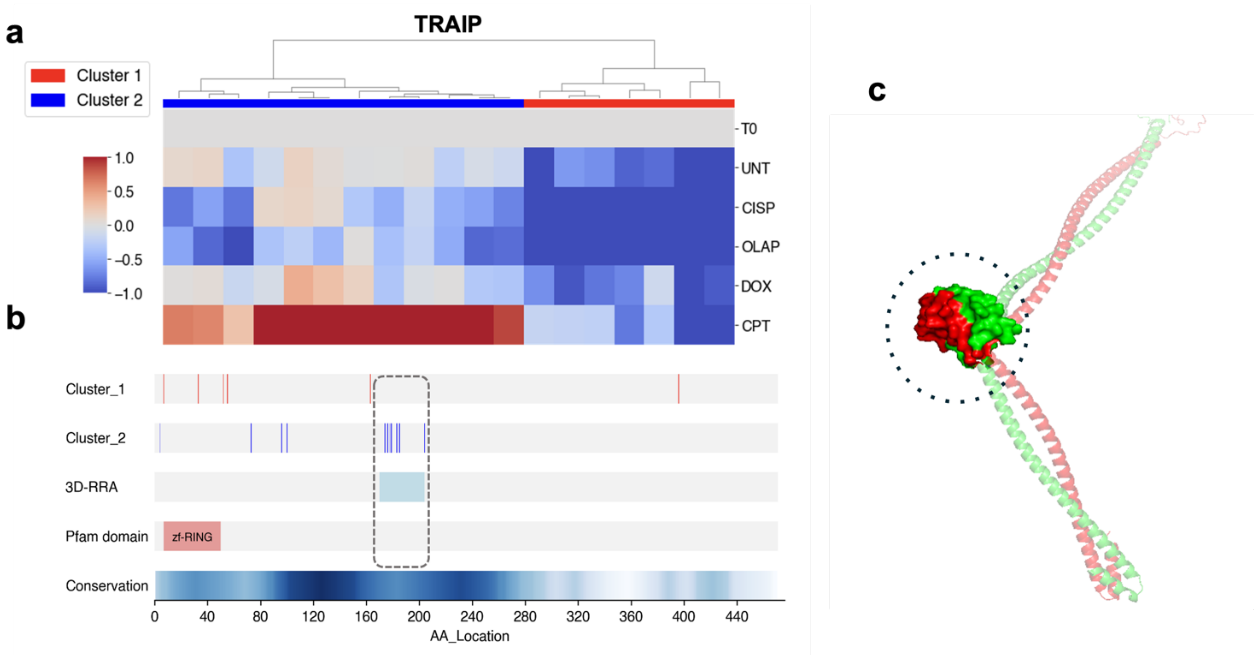


**Figure S3. "3D-RRA" analysis of TRAIP in DDR dataset.** (a) A heatmap showing the clustering of functional mutations of TRAIP based on the phenotypic effects under different drug treatment conditions (UNT: Untreated, CISP: Cisplatin, OLAP: Olaparib, DOX: Doxorubicin, CPT: Camptothecin). The phenotypic scores represent log fold-change (LFC) in cell viability relative to T0 (pre-selection). (b) Distribution of mutations in different clusters identified in (a) across the 1D protein sequence of TRAIP, aligned with "3D-RRA" identified substructures, the Pfam domain annotation and AlphaFold-predicted conservation score. The unannotated substructure (AA 170-210) identified by ProTiler is highlighted with a dashed rectangular box. (c) Structural model of TRAIP homodimer, with substructure identified in (b) shown in surface representation and encircled with dashed line, as predicted by Alphafold.

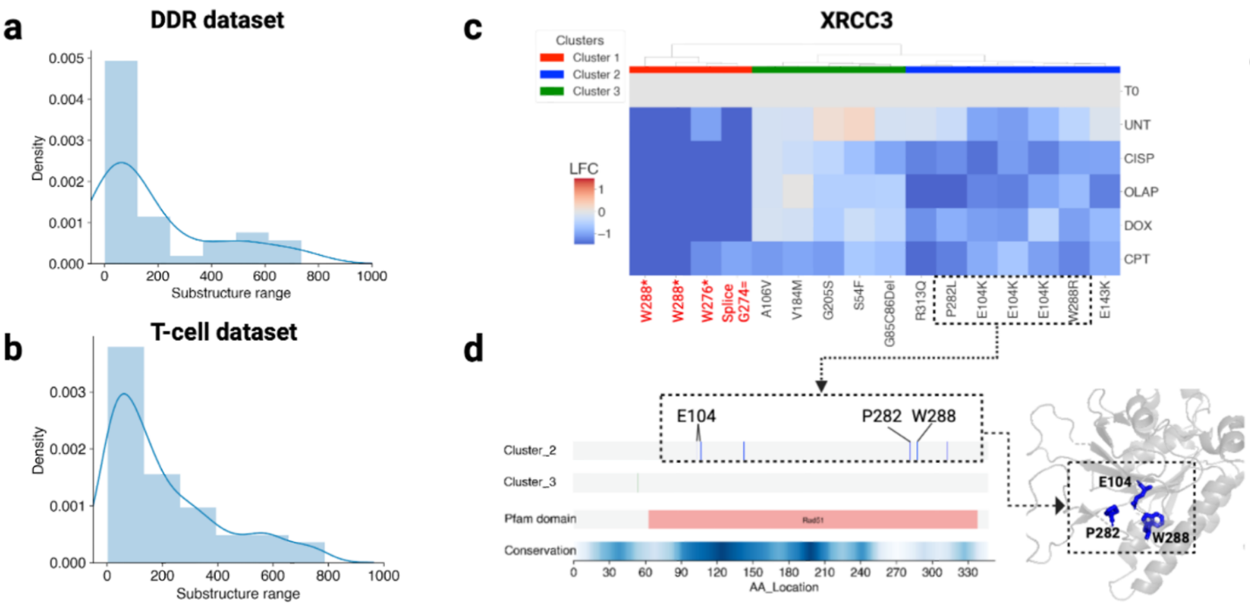


**Figure S4. ProTiler-Mut identifies long-range interplay among amino acid residues.** (a-b) Distribution of the maximum 1D distance among residues within ProTiler-Mut identified substructures in the (a) DDR dataset and (b) T-cell dataset. (c) Heatmap showing the clustering of functional mutations in XRCC3 in the DDR dataset based on phenotypic effects under different drug treatment conditions (UNT: Untreated, CISP: Cisplatin, OLAP: Olaparib, DOX: Doxorubicin, CPT: Camptothecin). The phenotypic scores represent log fold-change (LFC) in cell viability relative to T0 (pre-selection). (d) The distribution of mutations in different clusters identified in (c) across the 1D protein sequence of XRCC3, and three residues (E104,P282,W288) in cluster2 that are distant in 1D (left panel) but close in 3D (right panel) are highlighted with a dashed rectangular box.

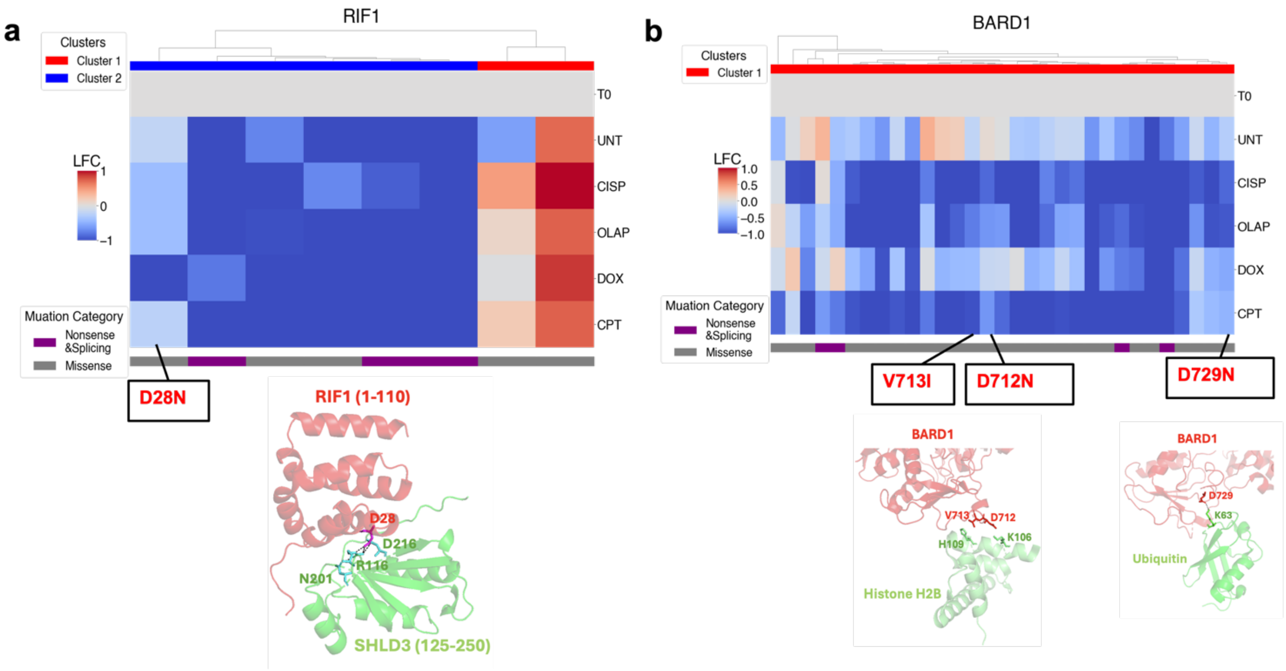


**Figure S5. ProTiler-Mut identifies well-characterized mutation-disrupted PPIs.** (a-b) Heatmaps (upper panel) showing the clustering of (a) RIF1 and (b) BARD1 in the DDR dataset based on the phenotypic effects under different drug treatment conditions (UNT: Untreated, CISP: Cisplatin, OLAP: Olaparib, DOX: Doxorubicin, CPT: Camptothecin). The phenotypic scores represent log fold-change (LFC) in cell viability relative to T0 (pre-selection). Color bars below the heatmap denote mutation types (purple for reference mutations, including nonsense and splicing; gray for missense). Mutations predicted to disrupt PPIs are highlighted in the rectangular box. The bottom panel displays the structural model of mutation-disrupted PPI interfaces of the (a) RIF1-SHLD3 complex and (b) BARD1-HistoneH2B and BARD1-Ubiquitin complexes.

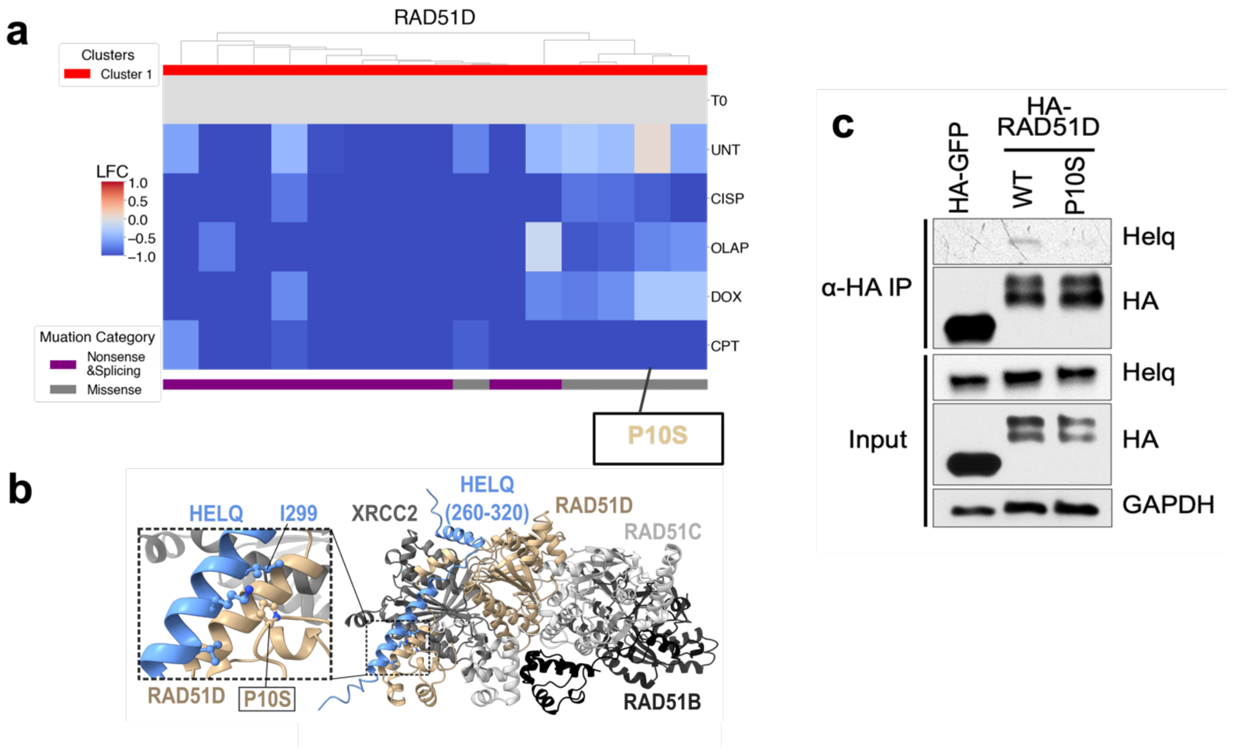


**Figure S6. PPI mapping and experimental validation for RAD51D functional mutations. (a)** Heatmap depicting clustering of functional mutations of RAD51D (rows) under different drug treatment conditions (UNT: Untreated, CISP: Cisplatin, OLAP: Olaparib, DOX: Doxorubicin and CPT: Camptothecin). The phenotypic scores represent log fold-change (LFC) in cell viability relative to T0 (pre-selection). Color bars below the heatmap denote mutation types (purple for reference mutations, including nonsense and splicing; gray for missense), the loss-of-function (LOF) mutation P10S, predicted to disrupt the RAD51Q–HELQ interaction, is highlighted with a rectangular box. **(b)** Structural model of the RAD51Q–HELQ complex, with a close-up view of the interaction interface. The LOF mutation P10S and its surrounding residues are shown in stick representation. **(c)** Co-immunoprecipitation (co-IP) assay showing the effect of the RAD51D P10S mutation on HEQL binding in HEK293T cells.

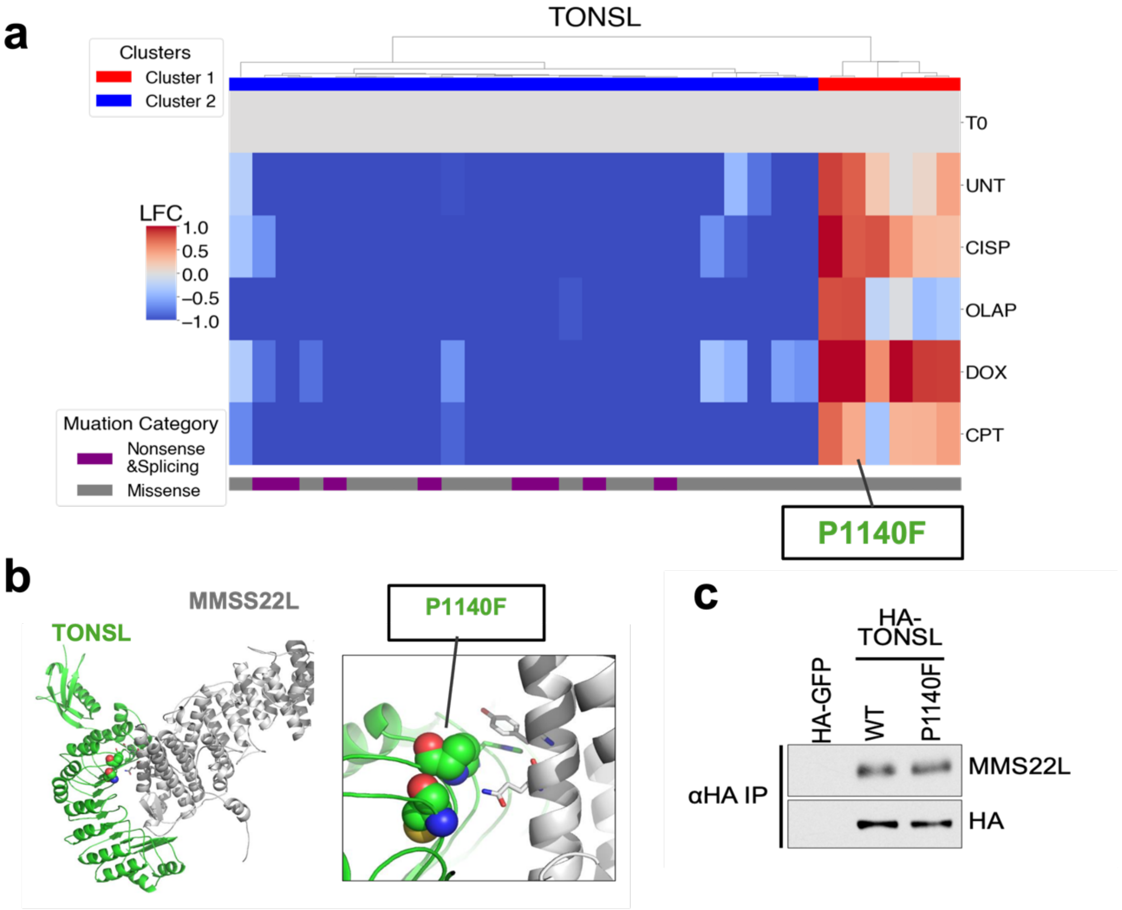


**Figure S7. PPI mapping and experimental validation of TONSL functional mutations. (a)** Heatmap depicting clustering of functional mutations of TONSL (rows) under different drug treatment conditions (UNT: Untreated, CISP: Cisplatin, OLAP: Olaparib, DOX: Doxorubicin and CPT: Camptothecin). The phenotypic scores represent log fold-change (LFC) in cell viability relative to T0 (pre-selection). Color bars below the heatmap denote mutation types (purple for reference mutations, including nonsense and splicing; gray for missense), the gain-of-function (GOF) mutation P1140F, predicted to disrupt the TONSL-MMSS22L interaction, is highlighted with a rectangular box. **(b)** Structural model of the TONSL-MMSS22L complex, with a close-up view of the interaction interface. The GOF mutation P1140F and its surrounding residues are shown in stick representation. **(c)** Co-immunoprecipitation (co-IP) assay showing the effect of the TONSL P1140F mutation on MMSS22L binding in HEK293T cells.

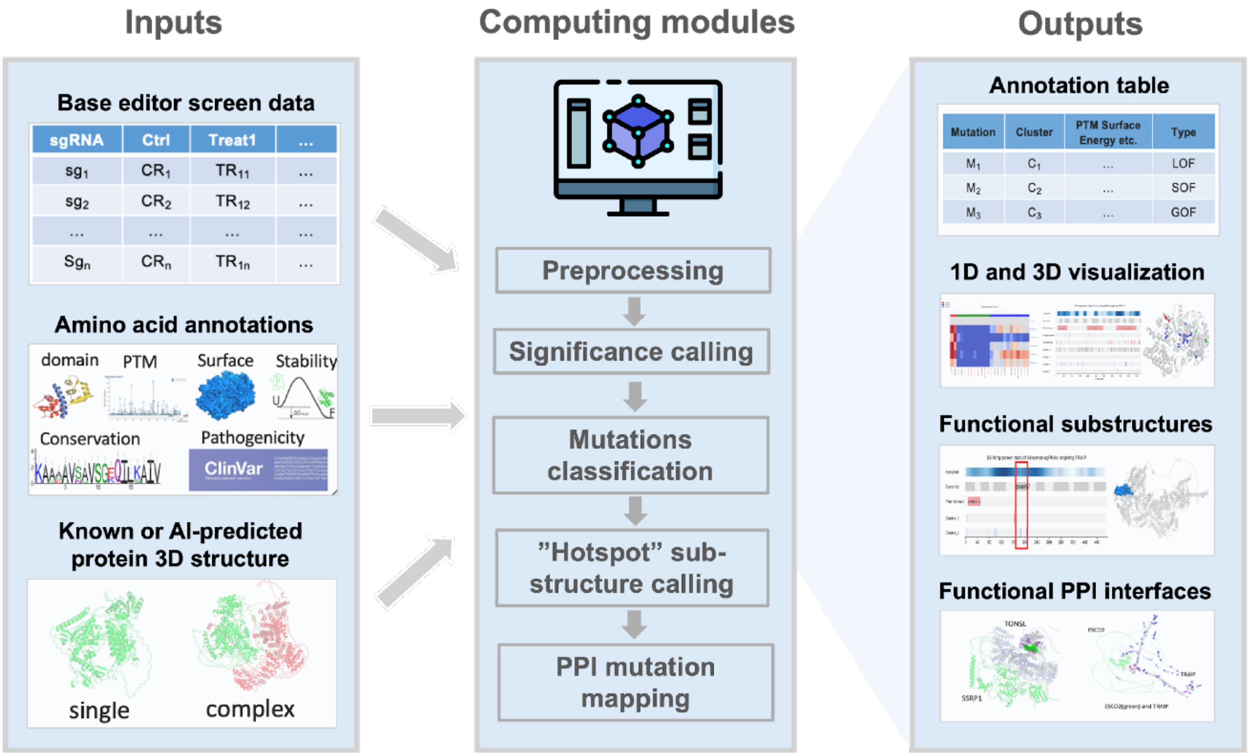


**Fig. S8. Schematic view of the ProTiler-Mut pipeline.** The pipeline begins with a read count table from tiling mutagenesis screens that includes detailed mutation data (amino acid position, change type, and functional annotation) along with their corresponding phenotypic effects under control (Ctrl) and treatment (Treat) conditions. Additional inputs include precise annotations of targeted residues and experimentally resolved or AI-predicted 3D protein structures, which may represent individual proteins or complexes. The pipeline involves several computational modules for streamlined analysis: 1). Preprocessing module for initial cleaning and preparation of the data; 2). Significance calling module for identification of functional mutations from the screening data; 3). Clustering module groups mutations based on their similarity and functional impact, further categorizing mutations into Loss-of-Function (LOF), Separation-of-Function (SOF), and Gain-of - Function (GOF) types; 4). The 3D RRA method for integrating spatial relationships and mutational effects to call significant substructures; 5). PPI-mapping module for systematically scanning all the experimental resolved or AI-predicted protein complexes to identify potential mutation-disrupted PPI interfaces for further experimental validation. The final outputs of the ProTiler-Mut pipeline includes a table with detailed mutation annotations, including mutation type, cluster information, PTM surface energy, and categorization; graphical representations of mutations and their effects in both linear (1D) and three-dimensional (3D) formats, as well as functionally significant protein substructures and mutation-disrupted PPI interfaces.

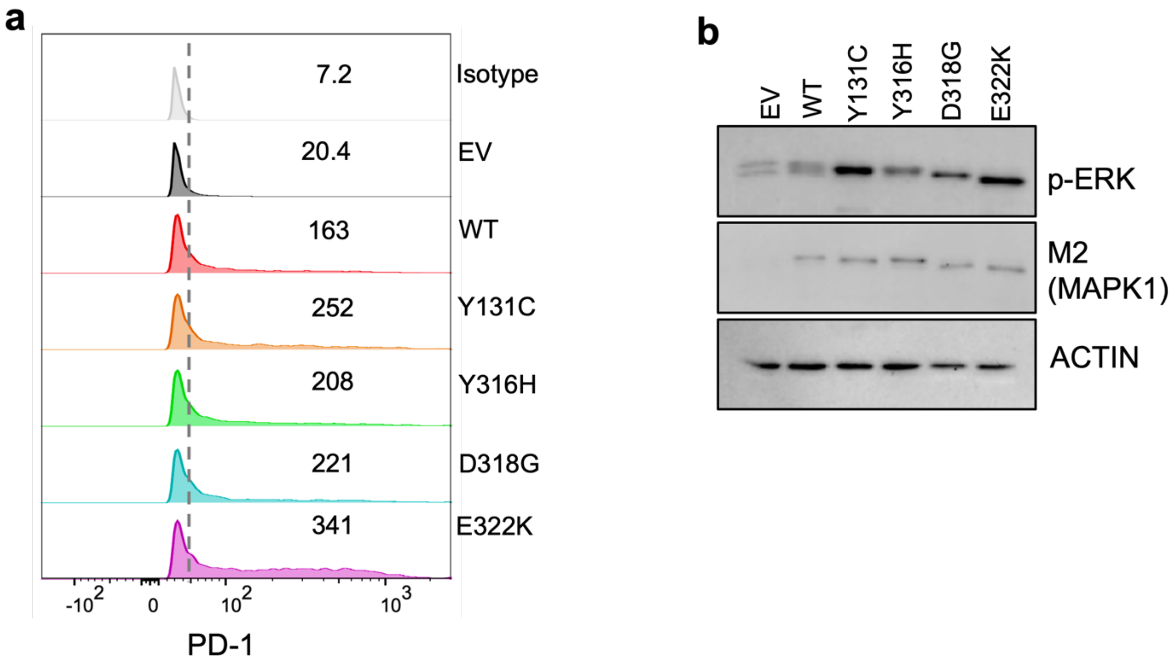


**Fig.S9. Analysis of PD-1 surface expression and MAPK1 phosphorylation in Jurkat cells.** **(a)** Jurkat cells were transfected with either empty vector (EV), wild-type (WT) or mutant MAPK1. PD-1 surface levels were assessed by FACS, and representative histograms from three independent experiments with mean fluorescence intensity (MFI) values are shown. **(b)** MAPK1 phosphorylation and expression of transfected constructs were evaluated by Western blot.

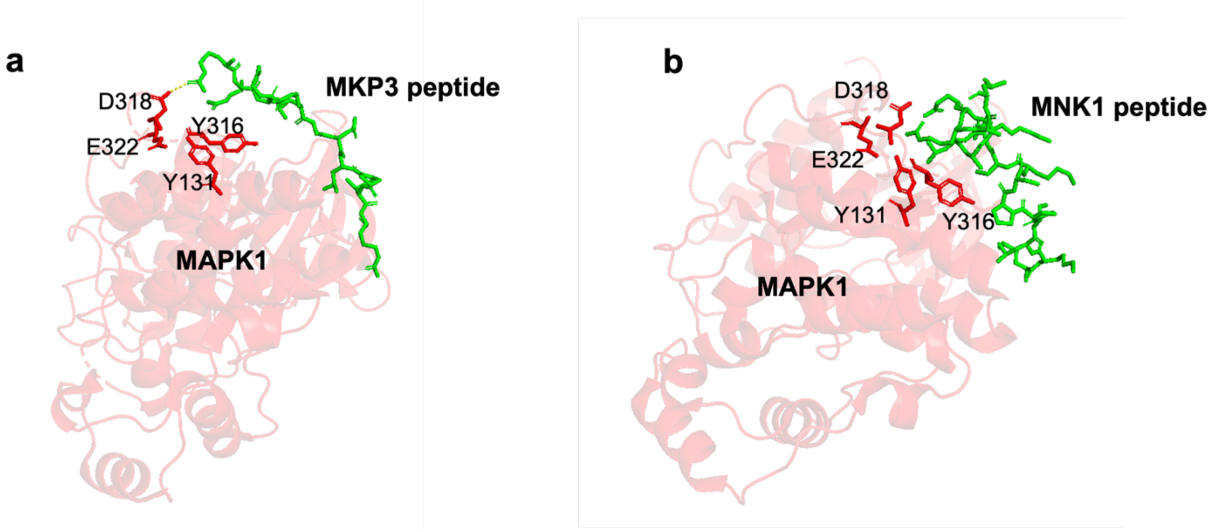


**Fig.S10. (a-b).** Structural model of MAPK1 (red) in complex with a linear motif in **(a)** MKP3 (green, PDB: 2FYS) and **(b)** MNK1 (green, PDB: 2Y9Q), with the interaction interface between MAPK1 substructure and peptide highlighted in sticks representation.

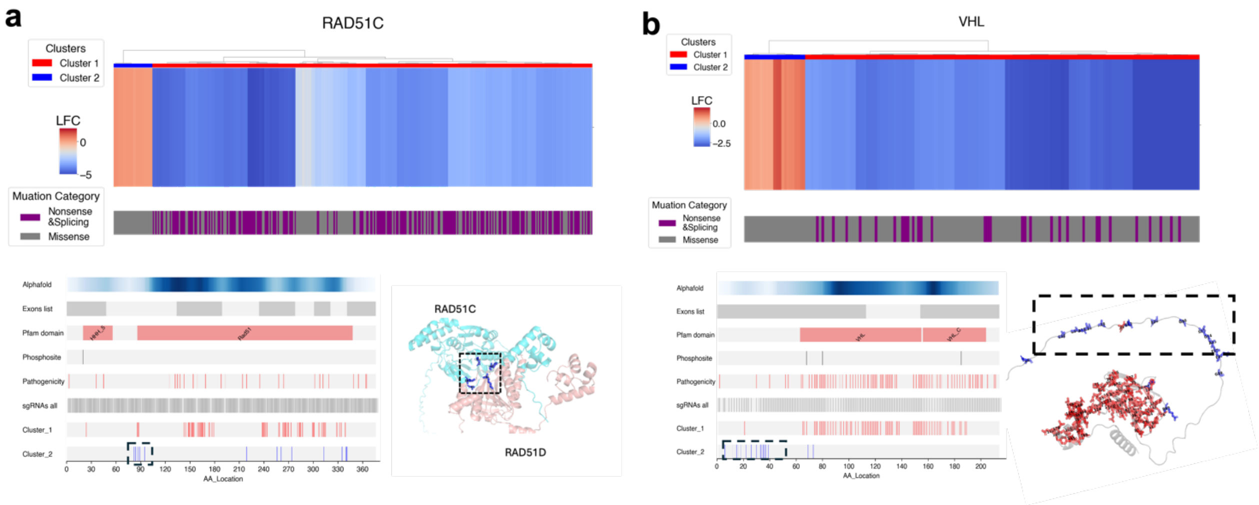


**Fig. S11. Application of ProTiler-Mut to HDR-mediated saturated genome editing (SGE) datasets.** (a-b) Analysis of SGE data targeting (a) RAD51C and (b) VHL. The upper panel shows the heatmaps depicting clustering of functional mutations based on their effects on cell viability, the color bar below the heatmap denote mutation types (purple for reference mutations, including nonsense and splicing; gray for missense). The bottom left panel shows the distribution of mutation in different clusters across the 1D protein sequence of the targeted protein, aligned with AlphaFold-predicted conservation scores, exons distributions, the Pfam domain annotation, the phosphorylation site annotations, as well as the pathogenicity annotation from ClinVar. The bottom right panel in (a) shows the 3D structure model of the RAD51C-RAD51D complex, highlighting the functional GOF substructures that are enriched in the interaction interface with dashed rectangular box. The bottom right panel in (b) shows the 3D structure model of VHL. The GOF cluster enriched at the N-terminal region is highlighted in a dashed box.
